# Interplay between mitochondria and reactive oxygen and nitrogen species in metabolic adaptation to hypoxia in facioscapulohumeral muscular dystrophy: potential therapeutic targets

**DOI:** 10.1101/2021.09.08.459509

**Authors:** P Heher, M Ganassi, A Weidinger, EN Engquist, J Pruller, TH Nguyen, A Tassin, AE Declèves, K Mamchaoui, J Grillari, AV Kozlov, PS Zammit

## Abstract

Facioscapulohumeral muscular dystrophy (FSHD) is characterised by descending skeletal muscle weakness and wasting. FSHD is caused by mis-expression of the transcription factor DUX4, which is linked to oxidative stress, a condition especially detrimental to skeletal muscle with its high metabolic activity and energy demands. Oxidative damage characterises FSHD and recent work suggests metabolic dysfunction and perturbed hypoxia signalling as novel pathomechanisms. However, redox biology of FSHD remains poorly understood, and integrating the complex dynamics of DUX4-induced metabolic changes is lacking.

Here we pinpoint the kinetic involvement of altered mitochondrial RONS metabolism and impaired mitochondrial function in aetiology of oxidative stress in FSHD. Transcriptomic analysis in FSHD muscle biopsies reveals strong enrichment for pathways involved in mitochondrial complex I assembly, nitrogen metabolism, oxidative stress response and hypoxia signalling. We found elevated ROS levels correlate with increases in steady-state mitochondrial membrane potential in FSHD myogenic cells. DUX4 triggers mitochondrial membrane polarisation prior to oxidative stress generation and apoptosis through mitochondrial ROS, and affects NO· bioavailability via mitochondrial peroxidation. We identify complex I as the primary target for DUX4-induced mitochondrial dysfunction, with strong correlation between complex I-linked respiration and cellular oxygenation/hypoxia signalling activity in environmental hypoxia. Thus, FSHD myogenesis is uniquely susceptible to hypoxia-induced oxidative stress as a consequence of metabolic mis-adaptation. Importantly, mitochondria-targeted antioxidants rescue FSHD pathology more effectively than conventional antioxidants, highlighting the central involvement of disturbed mitochondrial RONS metabolism. This work provides a pathomechanistic model by which DUX4-induced changes in oxidative metabolism impair muscle function in FSHD, amplified when metabolic adaptation to varying O_2_ tension is required.

**Highlights:**

- Transcriptomics data from FSHD muscle indicates enrichment for disturbed mitochondrial pathways
- Disturbed RONS metabolism correlates with mitochondrial membrane polarisation and myotube hypotrophy
- DUX4-induced changes in mitochondrial function precede oxidative stress through mitoROS and affect hypoxia signalling via complex I
- FSHD is sensitive to environmental hypoxia, which increases ROS levels in FSHD myotubes
- Hypotrophy in hypoxic FSHD myotubes is efficiently rescued with mitochondria-targeted antioxidants

**Graphical abstract:** 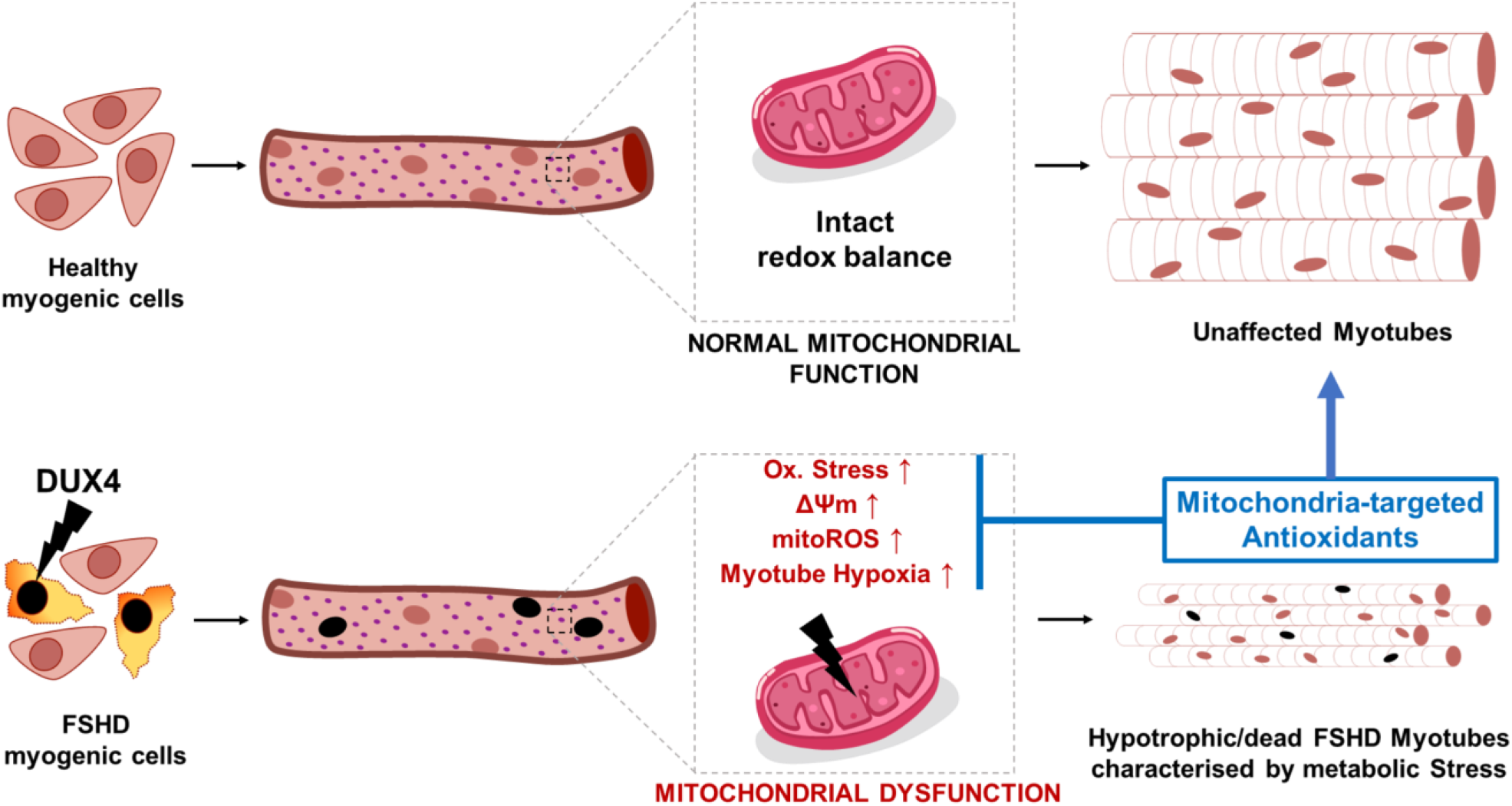

## INTRODUCTION

Facioscapulohumeral muscular dystrophy (FSHD; OMIM 158900) is an incurable, hereditary disease primarily affecting skeletal muscle. FSHD is the third most common inherited muscular dystrophy, with an estimated prevalence of 4-12 in 100000 [1, 2]. FSHD presents as a descending, often left/right asymmetric, skeletal muscle weakness and atrophy, starting in facial muscles such as orbicularis oculi and orbicularis oris, and progressing to muscles of the shoulder girdle/proximal arm, before affecting specific lower limb muscles [3, 4]. Fortunately, life-span is not attenuated, but FSHD severely affects quality of life, as approximately 30% of patients become wheelchair-bound while a further 10% require leg braces [3, 5]. In addition, some FSHD patients undergo fixation of the scapula to the ribcage to stabilise the shoulder, entailing incapacitation for months with associated muscle loss. The FSHD clinical phenotype also extends to extra-muscular features including high-frequency hearing loss that is often sensorineural and may progress to deafness [6], and/or retinal telangiectasia (Coat’s disease), a condition where abnormal vascular growth in the macula causes progressively compromised vision [7]. FSHD is a highly heterogeneous pathology, with presentations varying dramatically between first-degree relatives and even mono-zygotic twins [8, 9]. Furthermore, males and females exhibit differential penetrance, with males usually presenting in the second decade of life while females are typically affected by the third [10].

The genetic cause underlying FSHD pathogenesis is epigenetic derepression of the subtelomeric D4Z4 macrosatellite repeats on chromosome 4q35 [4, 11]. Each D4Z4 unit contains an open reading frame for the *Double Homeobox 4 (DUX4)* retrogene, encoding the homeobox germline transcription factor DUX4 (OMIM 606009) [12, 13]. Approximately 95% of cases (FSHD1) are associated with contraction from the usual 11-100 repeats to <10 and CpG/DNA hypomethylation of the D4Z4 repeat region in the subtelomere of “permissive” chromosome 4qA allelic variants, [14–16]. Along with reduced repressive histone modifications, hypomethylation results in DUX4-full length transcription from the otherwise somatically repressed distal-most D4Z4 unit [17]. A polymorphism in FSHD-permissive 4qA haplotypes (4qA161 and rarer 4qA159 and 4qA168) provides a polyadenylation signal (PAS) for *DUX4* transcripts, allowing their stabilisation and translation [18]. Residual number of D4Z4 units inversely correlates with disease severity but at least one unit is needed for FSHD pathology [19]. FSHD2 (FSHD2; OMIM 158901) accounts for the remaining 5% of cases, which are mostly characterized by epigenetic derepression at D4Z4 through mutations in *Structural maintenance of chromosomes flexible hinge domain containing 1 (SMCHD1)* [20], and much more rarely with mutations in *DNA methyltransferase 3B (DNMT3B)* [21] or the SMCHD1 protein interactor *ligand-dependent nuclear receptor-interacting factor 1 (LRIF1)* [22]. *SMCHD1* encodes a chromatin modifier that controls and maintains CpG methylation patterns for inheritable epigenetic silencing, for example during X-chromosome inactivation [23]. FSHD2 patients however, often have D4Z4 repeat numbers towards the shorter end of the normal range (<20 repeats) on permissive 4qA haplotypes, so that hypomethylation again permits *DUX4* expression from the distal-most D4Z4 unit [20, 21].

While the genetics of FSHD have been studied in detail, much less is known about pathomechanisms. One of the most drastic consequences of aberrant *DUX4* expression is apoptosis, possibly through the p53-p21 axis [17, 24, 25] but molecular pathways are elusive. Further, DUX4 interferes with myogenic differentiation by inducing a more stem-cell-like transcriptional program [26], so likely impinging both developmental and regenerative myogenesis. DUX4 leads to rapid down-regulation of the transcription factors Myogenic Differentiation (MyoD) and Myogenic Factor 5 (Myf5), two key muscle regulatory factors (MRFs) [27, 28]. *MyoD* is a Paired Box 7 (PAX7) target gene, and sequence similarity between the PAX7 and DUX4 homeodomains suggests that DUX4 interferes with the transcriptional circuitry controlled by PAX7 [28]. Indeed, a PAX7 target gene score is globally repressed in FSHD biopsies [29], with PAX7 target gene score repression a potent biomarker for FSHD [30].

Oxidative stress is a well-known pathomechanism of muscle diseases, with redox imbalances in several disorders such as Duchenne muscular dystrophy (DMD) [31, 32], myotonic dystrophy type 1 (DM1) [33] and limb-girdle muscular dystrophy (LGMD) [34, 35]. A pioneering study by Winokur et al. [36] revealed that FSHD myoblasts show a transcriptional dysregulation of the oxidative stress response, with specifically robust downregulation of antioxidant enzymes involved in the glutathione-based redox system. FSHD myoblasts have higher susceptibility to exogenously induced oxidative stress than controls, a susceptibility not seen in cellular models of other muscular dystrophies. Since this seminal observation of a unique oxidative stress-related pathomechanism in FSHD, oxidative stress/damage and mitochondrial dysfunction have been established as hallmarks of the disease. Several other *in vitro* studies have found perturbations of oxidative stress response and cellular bioenergetic pathways on the mRNA and protein level [37–42]. Although FSHD myoblasts are generally capable of repairing moderate oxidative damage, they fail to do so when oxidative stress becomes high and/or chronic [43]. It is, however, unclear whether reduced capacity of cellular antioxidant systems and/or chronically increased levels of reactive oxygen (RONS, or ROS) and nitrogen (RNS) species trigger oxidative damage in FSHD. FSHD myoblasts show elevated ROS levels and oxidative DNA damage, which can be reduced by antioxidant treatment [44]. Likewise, footprints of oxidative stress/damage have been identified in FSHD patients, both intramuscular (lipid peroxidation, lipofuscin accumulation, altered protein carbonylation) and systemic (reduced antioxidant levels in blood) [42]. DUX4 confers susceptibility to oxidative stress-induced cell death, and DUX4 increases ROS levels which are rescued by DUX4 knockdown or administration of antioxidants [28].

Skeletal muscle has high metabolic activity, so myofibers have to constantly adapt and respond to intrinsic and microenvironmental changes in their redox and bioenergetic regulatory pathways to meet energy demand. Since dynamic interplay between RONS, mitochondria and O_2_/hypoxia signalling is core to muscle metabolic adaptation, chronic insult to the fine balance between pro- and antioxidant redox-mechanisms will lead to metabolic stress [45, 46].

Mitochondrial dysfunction in FSHD pathogenesis is under studied. An interesting respirometric study on patient muscle biopsies by Turki et al. [42] identified reduced cytochrome C oxidase (COX) activity and adenosine triphosphate (ATP) production in FSHD muscles, along with a decreased ratio between reduced and oxidized glutathione (GSH/GSSG). Both systemic oxidative stress and mitochondrial dysfunction correlated with muscle functional impairment, emphasizing the central role of metabolic stress in FSHD. Further, different levels of proteins involved in mitochondrial oxidative metabolism have been found between healthy, DMD and FSHD muscle, most notably complex I subunits such as NADH dehydrogenase flavoprotein (NDUFV) and NADH-ubiquinone oxidoreductase (NDUFA), as well as the mitochondrial uncoupler adenine nucleotide translocator 1 (ANT1) [39]. Increased ANT1 correlates with enhanced ROS production and receptor of advanced glycation end products (RAGE) and nuclear factor kappa B (NF-κB) activity [40], suggesting mitochondrial involvement in pro-apoptotic signalling in FSHD muscle degeneration.

Both developmental and regenerative myogenesis are redox-sensitive and metabolic stress is a well-established negative regulator of myogenic differentiation [47]. *In vitro*, FSHD myotubes show morphologic features of aberrant differentiation, evident as a disorganized or hypotrophic phenotype [48, 49]. We have recently shown that differentiating FSHD muscle cells fail to fully activate a key mediator of mitochondrial biogenesis, the peroxisome proliferator-activated receptor gamma (PPARγ) coactivator 1α/estrogen-related receptor α (PGC1α/ERRα) axis, resulting in FSHD myotube hypotrophy [48]. PGC1α overexpression or treatment with ERRα agonists effectively rescues this FSHD hypotrophic phenotype, suggesting that mitochondrial (dys)function is central to impaired FSHD myogenesis. Since mitochondria are the main site of cellular O_2_ consumption, hypoxia signalling and metabolic adaptation to varying O_2_ availability might be directly affected by oxidative stress and dysfunctional FSHD mitochondria. We have previously also shown perturbed hypoxia-inducible factor 1α (HIF1α) signalling in FSHD muscle biopsies [29, 50], and a recent study identifies HIF1α signalling as the main driver of DUX4-induced muscle cell death [51].

Although unclear how altered muscle cell metabolism, oxidative stress and hypoxia signalling are involved in FSHD pathogenesis at the molecular level, antioxidant treatment has been proposed as a potential therapy for FSHD [52]. Antioxidants rescue aspects of DUX4 toxicity, such as oxidative DNA damage [44] and apoptosis [53], possibly by suppressing *DUX4* transcription itself, which increases under oxidative stress through a DNA damage response-dependent mechanism [54]. A clinical trial evaluating dietary antioxidant supplementation (vitamin C, vitamin E, zinc gluconate and selenomethionine) in FSHD patients demonstrated moderate muscle functional improvement with a concomitant alleviation of oxidative stress and damage [55, 56].

To date, only conventional, non-targeted antioxidants that mainly accumulate in the cytoplasm have been investigated in clinics. Since the respiratory chain can be a significant driver of ROS formation, especially in dysfunctional mitochondria, more targeted antioxidant-based therapeutic approaches might improve the so far rather moderate clinical outcomes. Indeed, a recent study [57] has shown that mitochondria-targeted antioxidant treatment in low level DUX4-inducible muscle cells can partially rescue aspects of DUX4-induced redox and differentiation defects, suggesting a role of mitochondrial ROS (mitoROS) in DUX4 toxicity.

The aim of this study is to understand how disturbed redox signalling and muscle metabolism integrate into current pathophysiologic paradigms to identify how mitochondria contribute to metabolic stress in FSHD. Our differential gene expression analysis of RNA sequencing (RNAseq) data of FSHD muscle biopsies [58] reveals strong enrichment of genes involved in processes related to mitochondria, with mitochondrial complex I (NADH dehydrogenase) assembly being the top differentially expressed gene ontology (GO) term. We identify that elevated mitochondrial membrane potential (ΔΨm) correlates with increased ROS levels in a panel of FSHD patient-derived muscle lines, which produces hypotrophic FSHD myotubes. DUX4 increases ΔΨm and ROS in a dose-dependent manner. Importantly, DUX4-induced changes in mitochondrial function and metabolic activity precede oxidative stress through mitochondrial ROS (mitoROS) formation, identifying mitochondria as the source of excess ROS. Respirometric analysis of DUX4-overexpressing muscle cells reveals that, in accordance with the transcriptomics data, complex I is the main target conferring mitochondrial dysfunction, while complex II-linked respiration is largely unaffected. DUX4 also differentially affects HIF1α stabilisation under environmental hypoxia in myoblasts versus myotubes, emphasising the relation between mitochondrial respiration and hypoxia signalling. HIF1α stabilisation correlates with complex I-linked respiration, in a manner involving both cellular redistribution of O_2_ and mitoROS. DUX4-induced perturbation of cellular respiration and hypoxia signalling sensitizes FSHD myogenesis to hypoxia through increased oxidative stress, evident by hypoxia-induced increases in ΔΨm and ROS levels that aggravate the hypotrophic myotube phenotype compared to normoxia. Finally, we show that antioxidant treatment of differentiating FSHD muscle cells rescues morphological FSHD phenotypes in hypoxia through alleviation of oxidative stress. Intriguingly, although all tested antioxidants rescue FSHD myotube hypotrophy, the mitochondria-targeted superoxide (O_2_^.-^) dismutase (SOD) mimetic mitoTEMPO is most efficient, not only reducing ROS levels but also ΔΨm and cellular hypoxia while increasing metabolic activity. This work suggests that oxidative stress and hypoxia signalling perturbation in FSHD are caused by mitochondrial dysfunction, and that mitochondria-targeted antioxidants may offer a novel therapeutic entry point to complement the current, more experimental therapies directed at reducing DUX4 levels.

## RESULTS

### Transcriptional deregulation of pathways involved in the mitochondrial respiratory chain in FSHD muscle biopsies

We have previously shown that genes involved in oxidative phosphorylation (OXPHOS), mitochondrial aerobic metabolism and mitochondrial biogenesis are dynamically repressed in FSHD myogenesis [48]. Here, to examine interplay between mitochondria, RONS metabolism and hypoxia signalling, we performed transcriptional analysis on RNAseq data from magnetic resonance imaging (MRI)-guided FSHD muscle biopsies [58]. We identified 7035 differentially expressed genes (DEG), with 3147 down-regulated and 3888 up-regulated in muscle biopsies from 6 FSHD patients with severe pathology, compared to 9 control individuals (Fig. 1A). GO analysis identified significantly enriched biological processes related to mitochondria, response to oxidative stress and O_2_ levels, and metabolism of nitrogen compounds (Fig. 1B, C), regulated through differential expression of 887 genes in FSHD (Fig. 1D). Notably, among the 30 top significant GO: biological processes (GOBPs), genes involved in mitochondrial complex I assembly and mitochondrial gene expression were specifically enriched. In addition, genes involved in development of functional muscle, blood vessels and immune response were found enriched in FSHD (Fig. 1B,C), identifying extra-muscular pathological features.

**Fig. 1:**
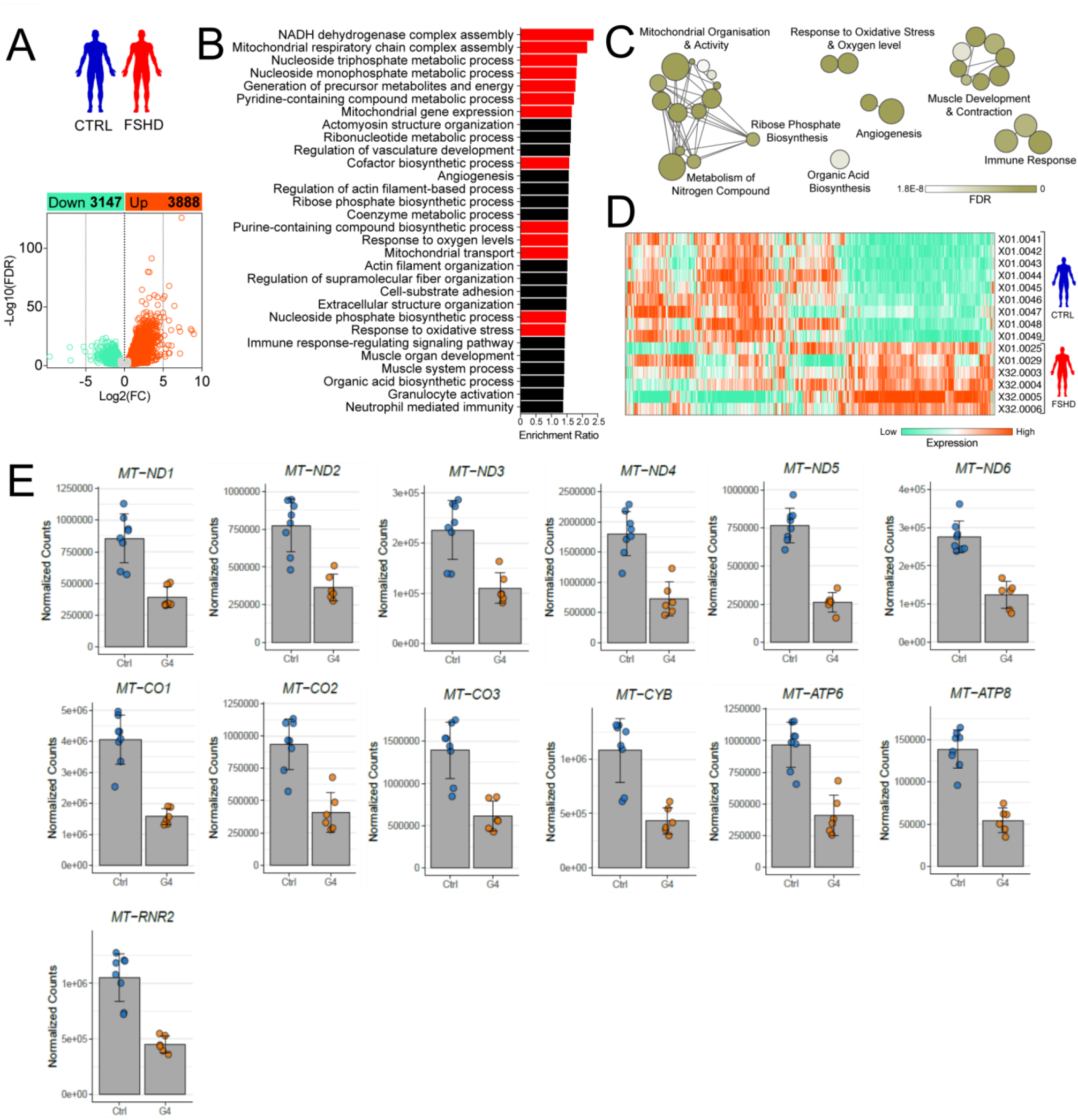
Transcriptional deregulation of pathways involved in mitochondrial oxidative metabolism, oxidative stress and hypoxia signalling in FSHD muscle biopsies. **(A)** 7035 genes were differentially expressed, with 3147 down-regulated and 3888 up-regulated in muscle biopsies from 6 FSHD patients with severe pathology, compared to 9 control individuals (GSE115650 from [58]). (**B, C**) Gene ontology analysis reveals significantly enriched biological processes related to mitochondria, response to oxidative stress and O_2_ levels and metabolism of nitrogen compound (marked in red), (**D**) regulated through differential expression of 887 genes in FSHD. (**E**) Robust transcriptional downregulation of all 13 protein-encoding genes of the mitochondrial genome.

Since the mitochondrial genome encodes important protein subunits involved in the respiratory chain [6 complex I subunits (MT-ND1-6), one complex III subunit (MT-CYB), three complex IV subunits (MT-CO1-3) and two complex V subunits (MT-ATP6, 8)], we also analysed expression of the 13 protein coding mitochondrial genes based on normalised counts from the same FSHD patients and unaffected individuals (Fig. 1E). In accordance with the GOBP analysis (Fig. 1B, C), all 13 mitochondrial genes were significantly downregulated in FSHD patient muscle. Since complex I subunits encoded by both the mitochondrial and nuclear genome are affected, mitochondrial dysfunction through alterations in complex I-linked respiration seems likely (Fig. 1B).

### FSHD myogenic cells are characterised by elevated ΔΨm and disturbed RONS metabolism

Since transcriptomic analysis showed widespread alterations in mitochondrial genes, we analysed the relationship between mitochondrial function characterised by ΔΨm and intracellular RONS metabolism in FSHD. ΔΨm is an important parameter of mitochondrial function, as is part of the proton motive force (pmf) maintained across the mitochondrial membrane to drive ATP synthesis through OXPHOS [59]. We examined ΔΨm in a panel of patient-derived FSHD myoblast cell lines: the 54 and K series derived from separate FSHD1 mosaic patients, and so isogenic bar the FSHD contraction, and 16s from an FSHD patient and their unaffected sibling. ΔΨm was significantly increased in FSHD muscle cells (54-12, K8 and 16A) compared to controls (54-6, K4, 16U), irrespective of stage of differentiation (myoblasts versus differentiated myotubes), using tetramethylrhodamine methyl ester (TMRM) fluorescence intensity measurements (Fig. 2A).

**Fig. 2:**
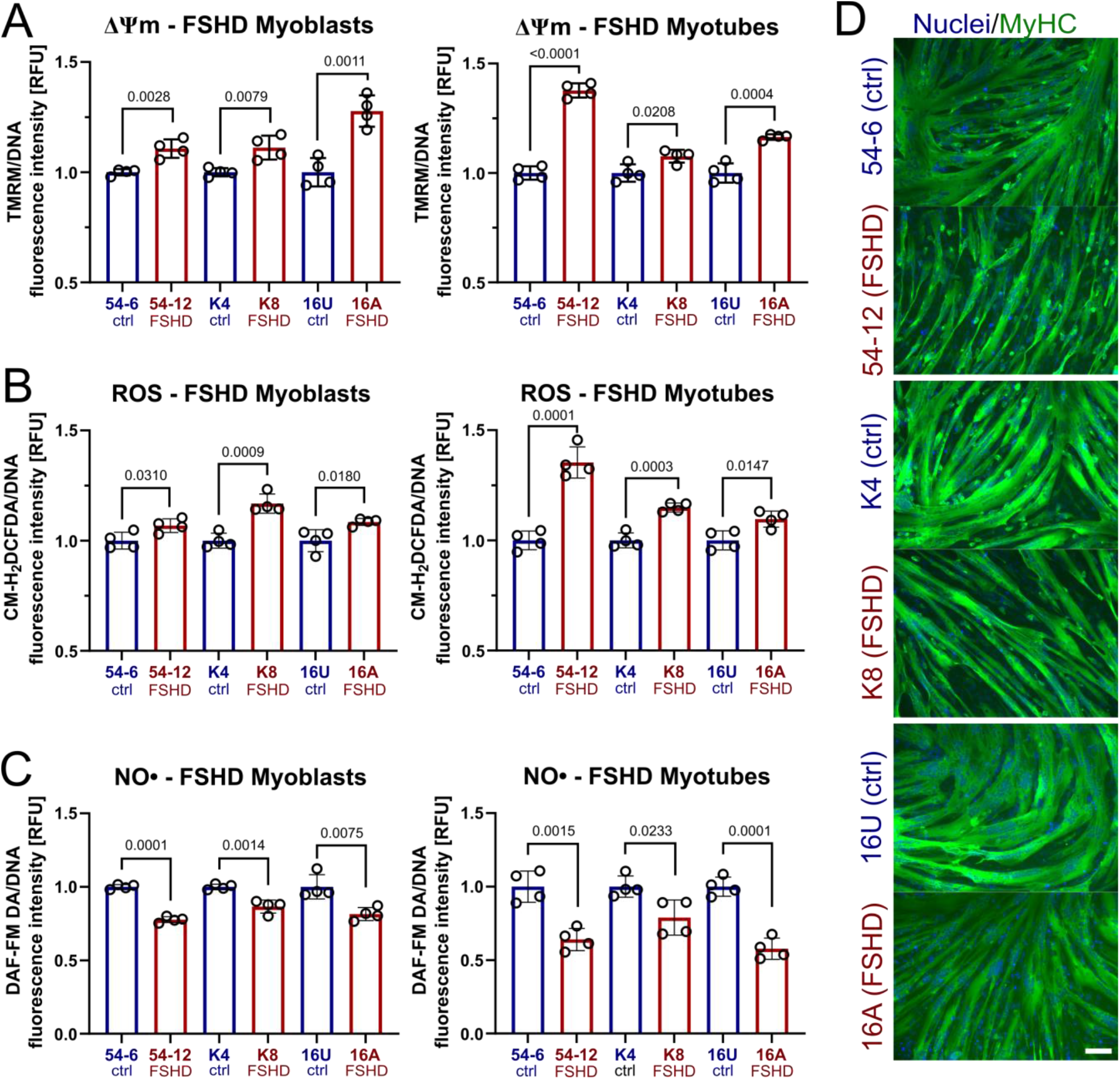
Altered RONS metabolism in FSHD muscle cells correlates with increased ΔΨm and produces hypotrophic myotubes. **(A)** Mitochondria in 3 independent FSHD myoblasts and myotube lines (54-6 ctrl/54-12 FSHD; K4 ctrl/K8 FSHD; 16U ctrl/16A FSHD) have consistently increased steady-state ΔΨm, along with **(B)** increased general (cytoplasmic) ROS and **(C)** decreased NO· levels. **(D)** Upon myogenic differentiation, FSHD myotubes exhibit a hypotrophic phenotype compared to their isogenic/sibling control (scale bar represents 100 μm). Data is mean ± s.d. from 3 independent cells pairs with 4 wells each from a representative experiment with *p* values as indicated.

Since the respiratory chain can produce large amounts of ROS at high ΔΨm [60], we next assessed general ROS and NO· levels using the general ROS indicator CM-H_2_DCFDA and the NO· probe DAF-FM DA. NO· was considered because it is known to facilitate mitoROS generation. We found consistently elevated ROS levels in FSHD myogenic cells (Fig. 2B), correlating with increased ΔΨm. Intriguingly, NO· levels were consistently reduced in FSHD myoblasts and, to a greater extent, in FSHD myotubes (Fig. 2C). This suggests peroxynitrite (ONOO^-^) formation through the reaction of mitochondrial O_2_ leaking from the electron transport chain (ETC) with NO· formed tentatively in the cytoplasm. These consistent changes in ΔΨm and RONS metabolism further correlated with formation of hypotrophic FSHD myotubes upon differentiation (Fig. 2D), supporting the notion that redox imbalances of a gradually more oxidative cellular environment, impair FSHD myogenesis.

### DUX4-induced changes in mitochondrial function and metabolic activity precede oxidative stress through increased mitoROS formation

To investigate the impact of DUX4 on the modulation of mitochondrial activity and RONS metabolism, we used the DUX4-inducible LHCN-M2-iDUX (iDUX4) human myoblast line [61] to examine the kinetics of redox changes in response to varying levels of DUX4. We titrated the doxycycline (DOX) inducer to elicit low (12.5 ng/mL DOX), medium (62.5 ng/mL DOX) and strong (125 ng/mL DOX) DUX4 induction. Similar to our observations in FSHD myoblasts, DUX4 induced an increase in ΔΨm in a dose-dependent manner (Fig. 3A), with changes in ΔΨm detectable after 8h with the highest DOX concentration (125 ng/mL) in iDUX4 myoblasts. After 12h, all DUX4 induction treatments showed significant elevation of ΔΨm, but ROS levels were not yet elevated. Metabolic activity at 12h was significantly reduced with all DUX4 expression regimes (Fig. S1A). A significant increase in ROS levels was then observed after 16h of medium and strong DUX4 induction. The minimum DUX4 induction that elicited a significant increase in ROS levels (62.5 ng/mL DOX) also yielded a significant increase in mitoROS levels at the 16h timepoint in iDUX4 myoblasts, as assessed with MitoTracker Red CM-H_2_XROS (Fig. 3B).

**Fig. 3:**
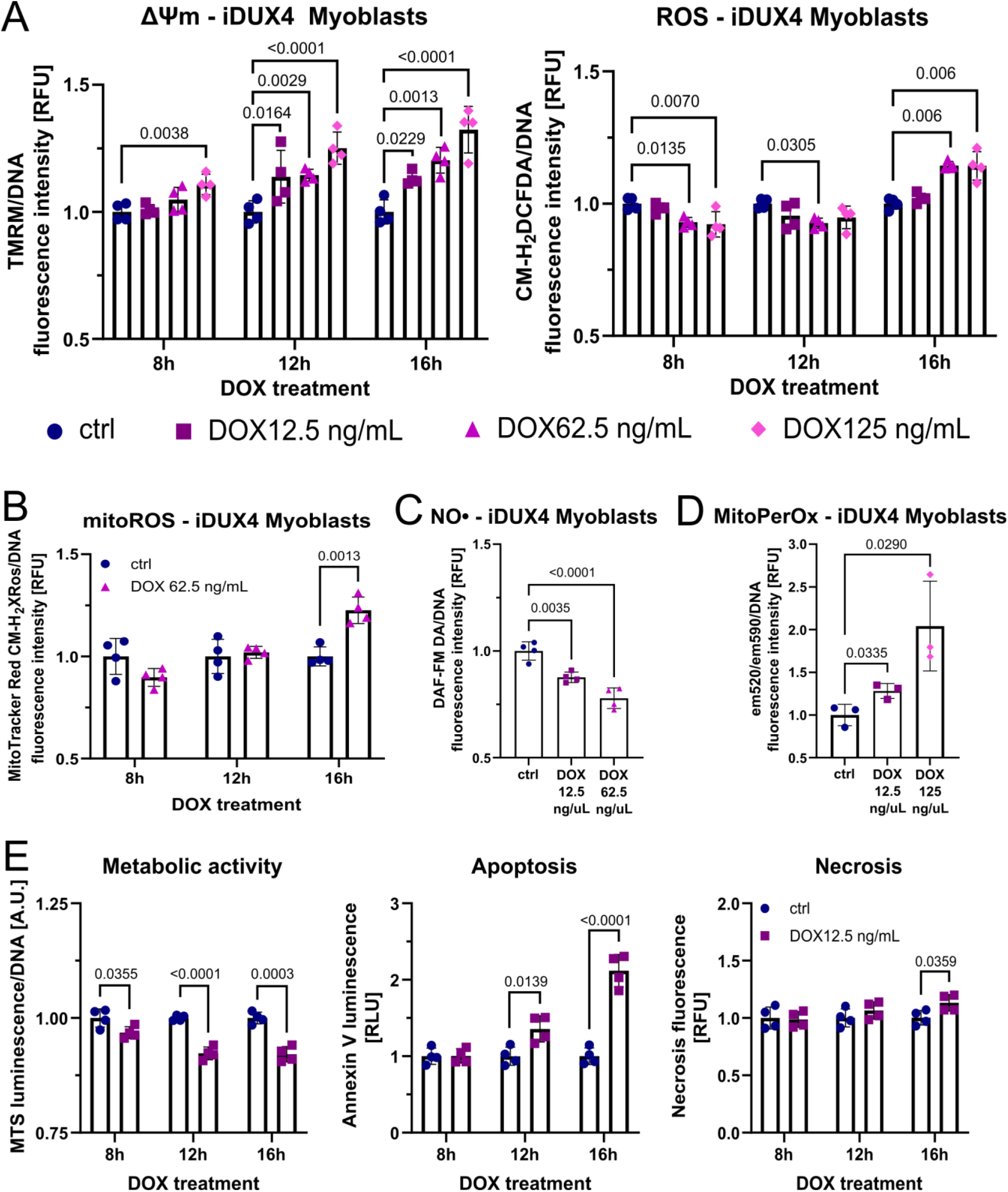
DUX4-induced changes in mitochondrial function are an early event in oxidative stress generation through mitochondrial ROS. **(A)** DUX4 increases ΔΨm in iDUX4 myoblasts in a dose dependent manner, preceding detection of elevated ROS levels by at least 4h, and **(B)** subsequently triggers oxidative stress through mitochondrial ROS. **(C)** The gradual depletion of NO· by mitoROS triggers ONOO^-^ formation, causing **(D)** mitochondrial oxidative damage through lipid peroxidation after 16h of DUX4 expression. **(E)** Changes in metabolic activity precede apoptosis, the main trigger of DUX4-induced muscle cell death, which manifests after 16h of DUX4 expression. Data is mean ± s.d. from 4 wells each from a representative experiment with *p* values as indicated.

Given the exponential relationship between ΔΨm and mitoROS generation, even moderate increases in ΔΨm can drastically increase ROS levels. Changes in ΔΨm preceded by at least 4 hours increases in general (cytoplasmic) ROS (Fig. 3A) and mitoROS (Fig. 3B) in iDUX4 myoblasts at medium DUX4 stimulation (62.5 ng/mL DOX). Similar to FSHD patient myoblasts, DUX4 also reduced NO· levels in a dose-dependent manner in iDUX4 myoblasts (Fig. 3C). Interestingly, moderate DUX4 induction (12.5 ng/mL DOX) did not lead to significant increase in ROS levels after 16h treatment but already affected NO· levels, possibly through ONOO^-^ formation within mitochondria (Fig. 3D), suggesting mitochondrially produced O_2_ is reducing bioavailability of intracellular NO· and subsequently causes mitochondrial oxidative damage as an early event of DUX4-mediated redox disturbance.

To investigate the relationship between redox balance and cell death, we used low DUX4 induction (12.5 ng/mL DOX) to track changes in metabolic activity and apoptosis/necrosis. DUX4 led to an immediate decrease in metabolic activity, already evident at the earliest 8h timepoint of DOX stimulation, with the first signs of apoptosis evident around 16h (Fig. 3E). Apoptosis seems to be the driving force of DUX4 toxicity, although a moderate increase in necrosis could also be detected. As expected, apoptosis and necrosis correlated with the degree of DUX4 induction (Fig. S1B, C), as did (mito)ROS levels and ΔΨm (Fig. 3A, B) in iDUX4 myoblasts. DUX4-induced apoptosis became significant for all DOX treatment groups after 16h and likely involves mitochondrial pro-apoptotic signalling, as onset of apoptosis was characterized by a subsequent gradual collapse of ΔΨm with a concomitant decrease in ROS levels, likely due to cell death (Fig. S1D).

To examine whether effects of DUX4 on ΔΨm and RONS metabolism are consistent between iDUX4 myoblasts and myotubes, we induced DUX4 expression at variable intensities in iDUX4 myotubes. We obtained similar results compared to myoblasts, but overall higher DOX concentrations were needed, as myotubes mainly rely on OXPHOS [62] and are thus generally better equipped to cope with resulting oxidative stress [63, 64]. Increase of ΔΨm in response to variable DUX4 expression was dose-dependent, with a strong induction (125 ng/mL DOX) leading to a significant increase after 12h, with elevated ROS levels evident after 24h of DOX treatment in iDUX4 myotubes (Fig. 4A). Likewise, this timepoint coincided with an increase in mitoROS levels (Fig. 4B), along with a reduction of NO· levels and ONOO^-^ formation already observed after low to medium DUX4 induction (Fig. 4C, D). All DOX treatment regimes lead to reduced metabolic activity after 24h in iDUX4 myotubes (Fig. 4E), pointing to similar mechanisms of DUX4-induced redox disturbances to those operating in iDUX4 myoblasts.

**Fig. 4:**
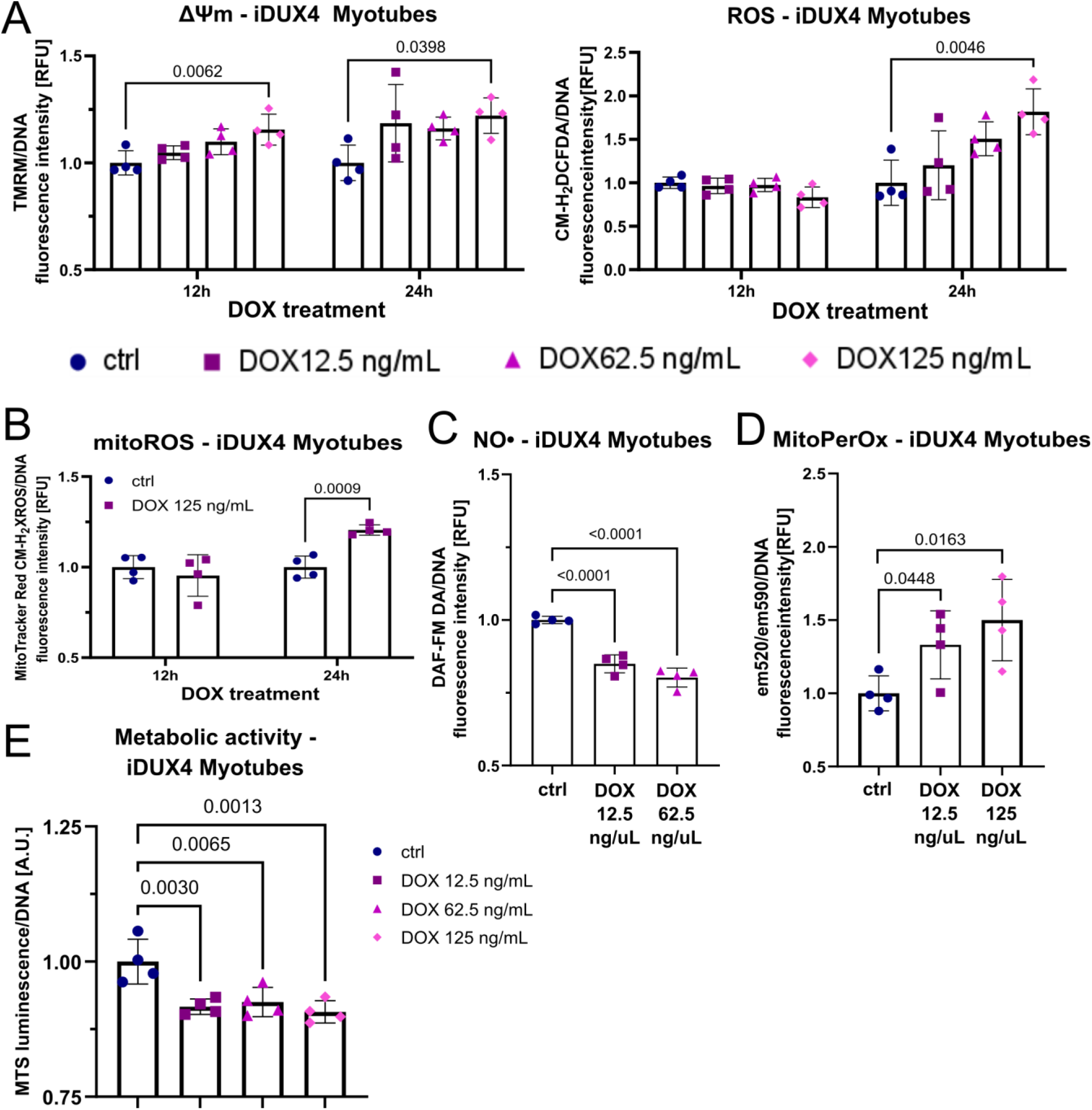
DUX4 expression in myotubes impacts mitochondrial function and subsequently perturbs RONS metabolism. **(A)** DUX4 expression increases ΔΨm in iDUX4 myotubes in a dose dependent manner, preceding detection of elevated ROS levels by 12h, and subsequently triggers **(B)** oxidative stress through mitochondrial ROS. **(C)** Dose-dependent reduction of NO· through mitoROS is accompanied by **(D)** mitochondrial oxidative damage after 24h of DUX4 expression. **(E)** Similar to changes in ΔΨm, DUX4 expression for 12h in myotubes causes reduction of metabolic activity before oxidative stress through elevated ROS becomes evident. Data is mean ± s.d. from 4 wells each from a representative experiment with *p* values as indicated.

### DUX4-induced mitochondrial dysfunction is conferred through mitochondrial complex I and interferes with hypoxia signalling

Having identified DUX4 as a major trigger of changes in ΔΨm and mitochondrial RONS metabolism, and the similarity of these changes between FSHD patient-derived and iDUX4 disease models, we next investigated effects of DUX4 on mitochondrial oxygen consumption through high-resolution respirometry. Normally, complex I contributes to generation of ΔΨm, so increased ΔΨm in FSHD might be attributable to altered complex I activity, as suggested by our transcriptomic analysis (Fig. 1). Thus, we focussed on how DUX4 affects mitochondrial respiration if electrons to the ETC are provided via either complex I or II.

We induced DUX4 in iDUX4 myoblasts with 62.5 ng/mL DOX for 16h, a concentration that robustly increases ΔΨm and (mito)ROS levels (Fig. 3A, B). DUX4 significantly reduced complex I-linked OXPHOS (state 3) and maximum electron transfer system (ETS) capacity in iDUX4 myoblasts, while complex II-linked respiration was unaffected (Fig. 5A-C). LEAK respiration (state 4) via complex I was also reduced (Fig. 5D), possibly contributing to mitoROS generation through mitochondrial membrane hyperpolarisation. LEAK respiration via complex II was unchanged (Fig. S2A), as were the respiratory control ratio (RCR) of both complex I and II (Fig. S2B). Representative oxygraphs are shown (Fig. S2C). Complex I-linked respiration was primarily affected by DUX4 expression in iDUX4 myotubes with 62.5 ng/mL DOX for 24h. However, DUX4 expression caused an increase in both complex I-linked OXPHOS (state 3) and maximum ETS capacity in iDUX4 myotubes, while these parameters where largely unchanged via complex II (Fig. 5E-G). In contrast to iDUX4 myoblasts, LEAK respiration via complex I (state 4) was unaffected (Fig. 4H), as was LEAK respiration through complex II (Fig. S2D). RCR of both complex I and II was moderately increased (Fig. S2E). Representative oxygraphs are shown (Fig. S2F). Myoblast viability and DNA content of sister myotube cultures were used for normalisation (Fig. S2G).

**Fig. 5:**
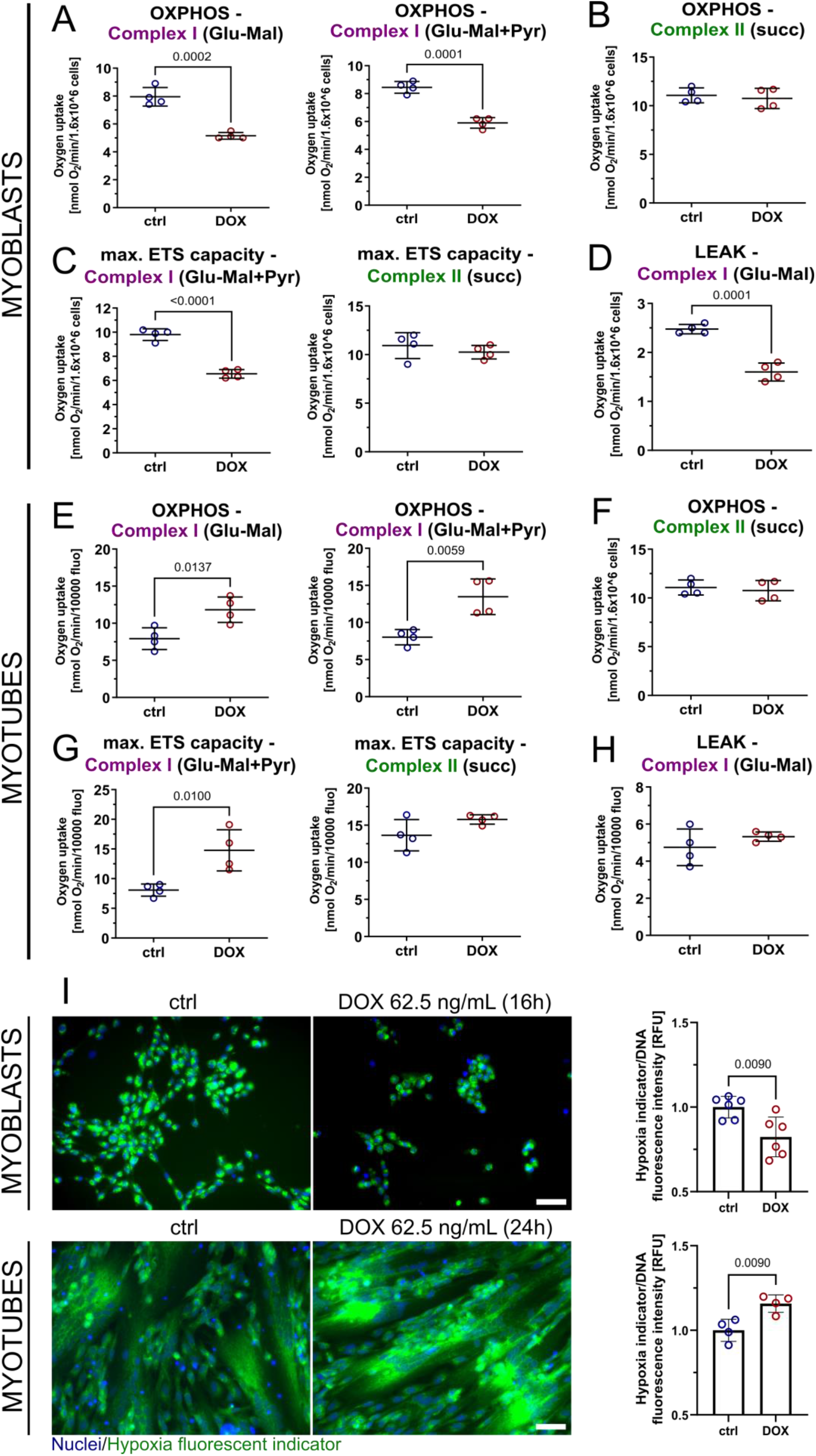
DUX4 affects mitochondrial respiration specifically at complex I and impairs cellular oxygenation through cellular redistribution of O_2_. **(A-D)** High-resolution respirometry in DUX4 expressing iDUX4 myoblasts (DOX 62.5 ng/mL for 16h) identifies reduced OXPHOS and LEAK (uncoupled) respiration through complex I, but not complex II. **(E-G)** Complex I-linked respiration is increased in DUX4 expressing myotubes (DOX 62.5 ng/mL for 24h), while complex II is again unaffected. **(H)** In contrast to myoblasts, DUX4 does not change LEAK respiration in iDUX4 myotubes. **(I)** Hypoxia indicator fluorescence microscopy of DUX4 expressing iDUX4 myoblasts (top panel) and myotubes (bottom panel) grown in hypoxia (1% O_2_) reveals correlation between complex I-linked respiration and cellular hypoxia, as quantified by indicator dye fluorescence intensity on a plate reader in a separate experiment (representative micrographs are shown, scale bar represents 50 μm). Data is mean ± s.d. from 4-6 wells each from a representative experiment with *p* values as indicated.

We next focused on the link between mitochondrial dysfunction and deregulated hypoxia signalling [50, 51]. Dysfunctional mitochondria with altered oxygen consumption rates can directly affect HIF1α nuclear stabilisation through mechanisms including redistribution of cellular O_2_ and RONS signalling [65–67]. We cultured iDUX4 myoblasts and myotubes in environmental hypoxia (1% O_2_) and induced DUX4 as per the respirometry experiments. Altered mitochondrial function correlated with cellular oxygenation: DUX4 expressing myoblasts (with reduced complex I-linked respiration) were marked by higher intracellular O_2_ levels (and were thus less hypoxic), whereas myotubes (with increased complex I-linked respiration) displayed lower intracellular O_2_ levels (and were thus more hypoxic), as assessed with fluorescence intensity measurements using a hypoxia sensitive fluorescent dye (Fig. 5I).

Differential regulation of the molecular response to hypoxia was further supported by the finding that DUX4 and HIF1α protein inversely correlated in myoblasts. Increasing DOX concentrations (12.5 vs 62.5 vs 125 ng/mL DOX for 24h) reduced HIF1α-positive nuclei from ∼100% in uninduced hypoxic iDUX4 myoblasts to ∼60% at the highest DUX4 levels, as assessed by immunofluorescence (Fig. 6A, B). Conversely, DUX4 induction in hypoxic iDUX4 myotubes increased the percentage of HIF1α-positive myonuclei from about 40% in uninduced controls to 60% at the highest DOX levels (Fig. 6C, D). Low to medium DUX4 induction in iDUX4 myotubes also led to significant changes in nuclear HIF1α in a dose-dependent manner, suggesting that DUX4 acts on HIF1α signalling activity in a mechanism involving mitochondria.

**Fig. 6:**
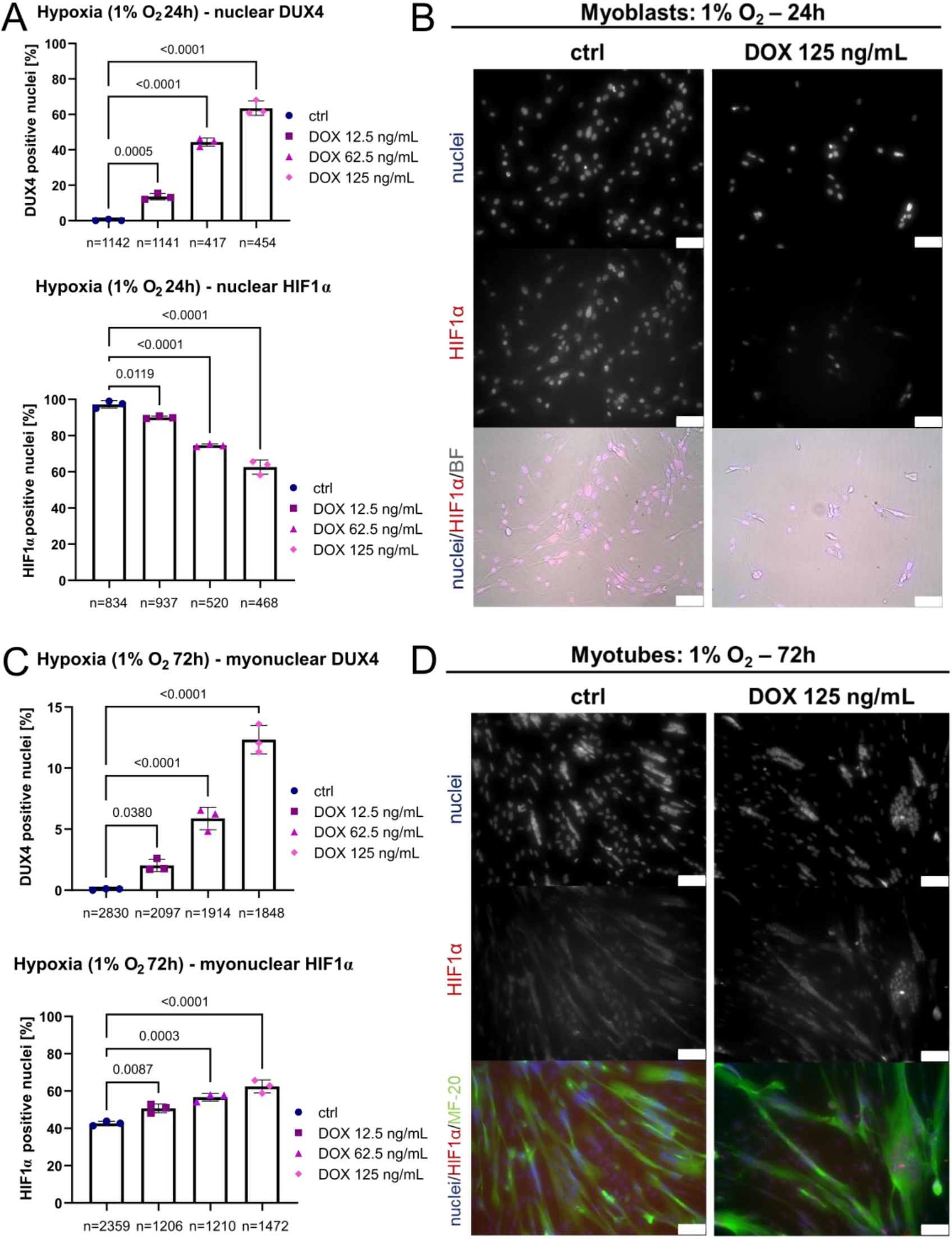
DUX4 interferes with HIF1a signalling activity in environmental hypoxia. **(A)** Percentage of DUX4- and HIF1α-positive nuclei in hypoxic iDUX4 myoblasts induced to express DUX4 for 24h at variable levels correlate inversely, as DUX4 expressing cells show reduced nuclear HIF1α (correlating with reduced complex I-linked OXPHOS; see Fig. 5A-D). **(B)** Representative immunofluorescence microscopy image of iDUX4 myoblasts after DUX4 expression (DOX 125 ng/mL for 24h) in hypoxia, compared to non-induced controls (scale bar represents 75 μm). **(C)** Percentage of DUX4-positive myonuclei in hypoxic myotubes induced to express DUX4 for 24h at variable amounts correlates with HIF1α nuclear localisation (and with increased complex I-linked OXPHOS; see Fig. 5E-G). **(D)** Representative immunofluorescence microscopy image of iDUX4 myotubes after DUX4 expression (DOX 125 ng/mL for 24h) in hypoxia, compared to non-induced controls (scale bar represents 75 μm). Data is mean ± s.d. of number of nuclei/myonuclei stated, from 3 wells from a representative experiment with *p* values as indicated.

### DUX4 aggravates hypoxia-induced oxidative stress and sensitises FSHD myogenesis to hypoxia

Having shown that DUX4 alters mitochondrial function and interferes with HIF1α activation, we next investigated the effects of DUX4 on ΔΨm and ROS levels in hypoxia. Myogenic differentiation is characterised by a gradual switch from predominantly glycolytic metabolism in myoblasts to OXPHOS in myotubes as the major energy source, and perturbation of metabolic pathways required for oxidative metabolism impacts on myogenic differentiation [68, 69]. Skeletal muscles are subject to constant physiological variations in local O_2_ availability, and hypoxia signalling is central to metabolic adaptation. Thus DUX4-induced redox and metabolic changes will interfere with this adaptation to hypoxia, a condition that naturally elicits some degree of oxidative stress [70].

We analysed ΔΨm and ROS levels in iDUX4 myoblasts and myotubes cultured in normoxia (21% O_2_) versus hypoxia (1% O_2_). Hypoxia increased both ΔΨm and ROS levels in non-induced iDUX4 controls. DUX4 increased ΔΨm and ROS levels in normoxia, and they were further enhanced at 1% O_2_ (Fig. 7A, B). Consequently, myogenic differentiation was affected at lower levels of DUX4 induction in hypoxia compared to normoxia. Non-induced control iDUX4 myoblasts differentiated into myotubes with similar efficiency regardless of the O_2_ availability at 21% versus 1%. However, myogenesis was impaired at lower DOX concentrations in hypoxia (62.5 ng/mL DOX) compared to normoxia (125 ng/mL DOX) as assessed by quantification of the myosin heavy chain (MHC) positive area (Fig. 7C, D). While high DUX4 induction with 125 ng/mL DOX elicited myotube hypotrophy and death both under normoxic and hypoxic conditions after 24 hours, medium induction (62.5 ng/mL DOX) only produced a phenotype in hypoxia (Fig. 7D). Morphologically, this phenotype was accompanied by iDUX4 myotube fragmentation and increased presence of hypotrophic myotubes with myonuclei strongly positive for HIF1α (Fig. S3). Morphological myotube deficits correlated with increased mitoROS levels, which were only evident at DOX concentrations that produced a hypotrophic phenotype (Fig. 7E).

**Fig. 7:**
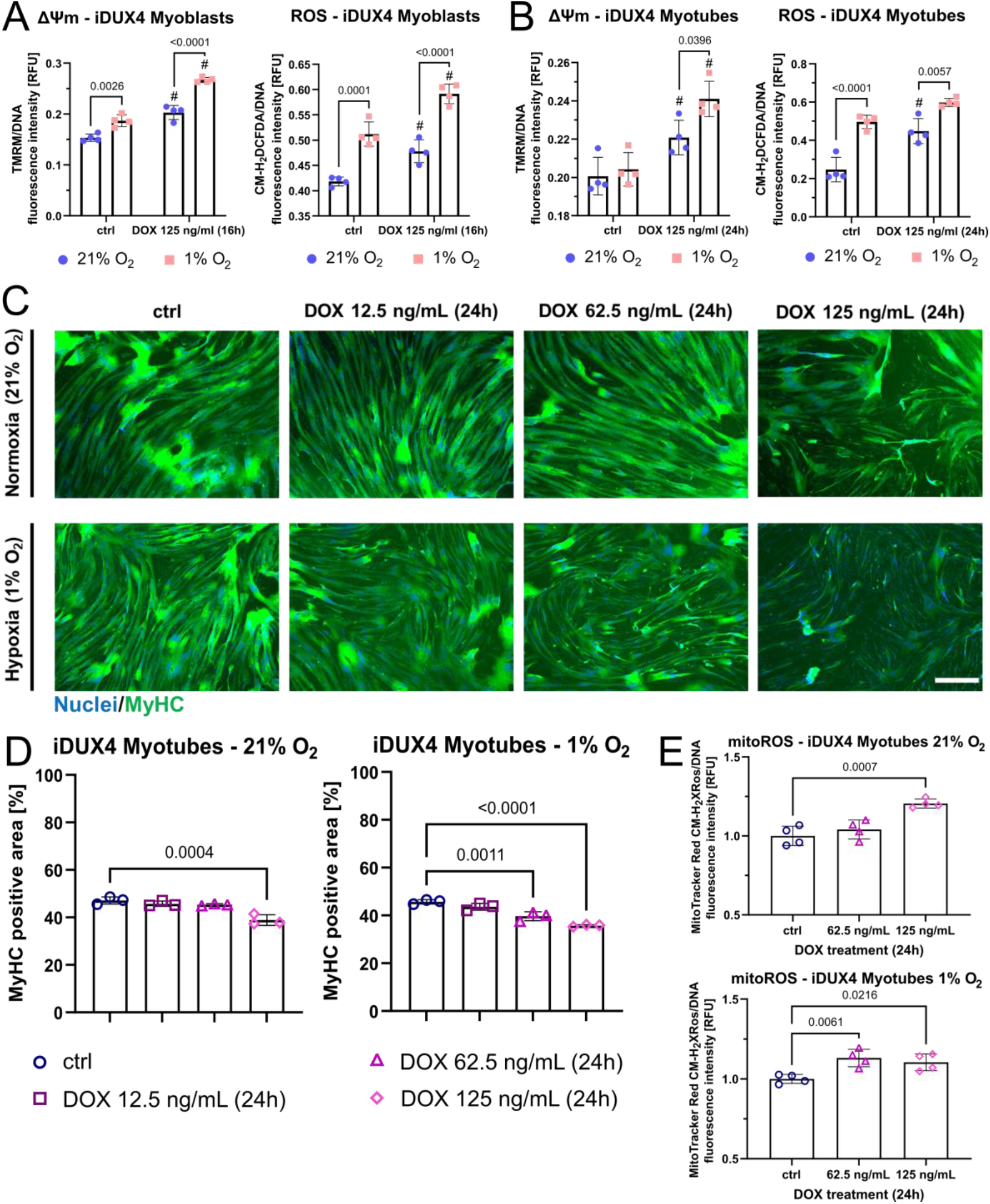
DUX4-induced mitochondrial dysfunction impairs myogenesis in hypoxia through aggravation of oxidative stress. **(A, B)** Hypoxia increases oxidative stress in non DUX4-induced iDUX4 myoblasts and myotubes. DUX4 expression increases ROS levels and ΔΨm regardless of O_2_ tension (# denotes significance between DUX4 induction at given O_2_ tension through DOX and the respective non-induced control), with ΔΨm increasing disproportionally in hypoxia. **(C, D)** Titration of the DUX4-inducer DOX (for 24h at variable amounts) in iDUX4 myotubes demonstrates that lower DUX4 levels are needed to produce a hypotrophic myotube phenotype in hypoxia compared to normoxia, as assessed by quantitation of the MyHC positive area from immunofluorescence micrographs (scale bar represents 500 μm). **(E)** DUX4 expressing myotubes are characterised by significantly elevated mitochondrial ROS levels at the minimal DOX concentration needed to elicit a phenotype in hypoxia versus normoxia. Data is mean ± s.d. from 3-4 wells each from a representative

We next assessed the effects of hypoxia on the three independent FSHD/control paired myoblast lines. FSHD myotubes maintained higher ΔΨm and ROS levels in hypoxia (Fig. 8A, B), with more pronounced differences from controls compared to normoxia (Fig. 2A, B). Since myotubes rely predominantly on OXPHOS, DUX4-induced interference with hypoxic adaptation should render FSHD myotubes more vulnerable to redox imbalances than FSHD myoblasts. We thus compared ROS levels of control versus FSHD myoblasts and myotubes between normoxia and hypoxia. Hypoxia did not elicit differential changes in ROS levels in FSHD and control myoblasts, but FSHD myotubes displayed a bigger increase in ROS levels compared to control myotubes when differentiated in hypoxia (Fig. 8C). ROS levels in hypoxic FSHD myotubes increased dramatically (approximately two-fold) compared to normoxia, and the concomitant increase of ΔΨm in FSHD myotubes (Fig. 8A) again suggests that this differential increase is caused by altered mitochondrial RONS metabolism. Notably, hypoxia consistently aggravated the hypotrophic FSHD myotube phenotype observed in normoxia in all three patient-derived models, whereas control myotubes did not show any gross morphological impairment of differentiation in hypoxia (Fig. 8D, E). This observation shows that the failure of FSHD myotubes to adapt their metabolism to hypoxia to maintain redox control and prevent oxidative stress/damage is the driving mechanism underlying myotube hypotrophy.

**Fig. 8:**
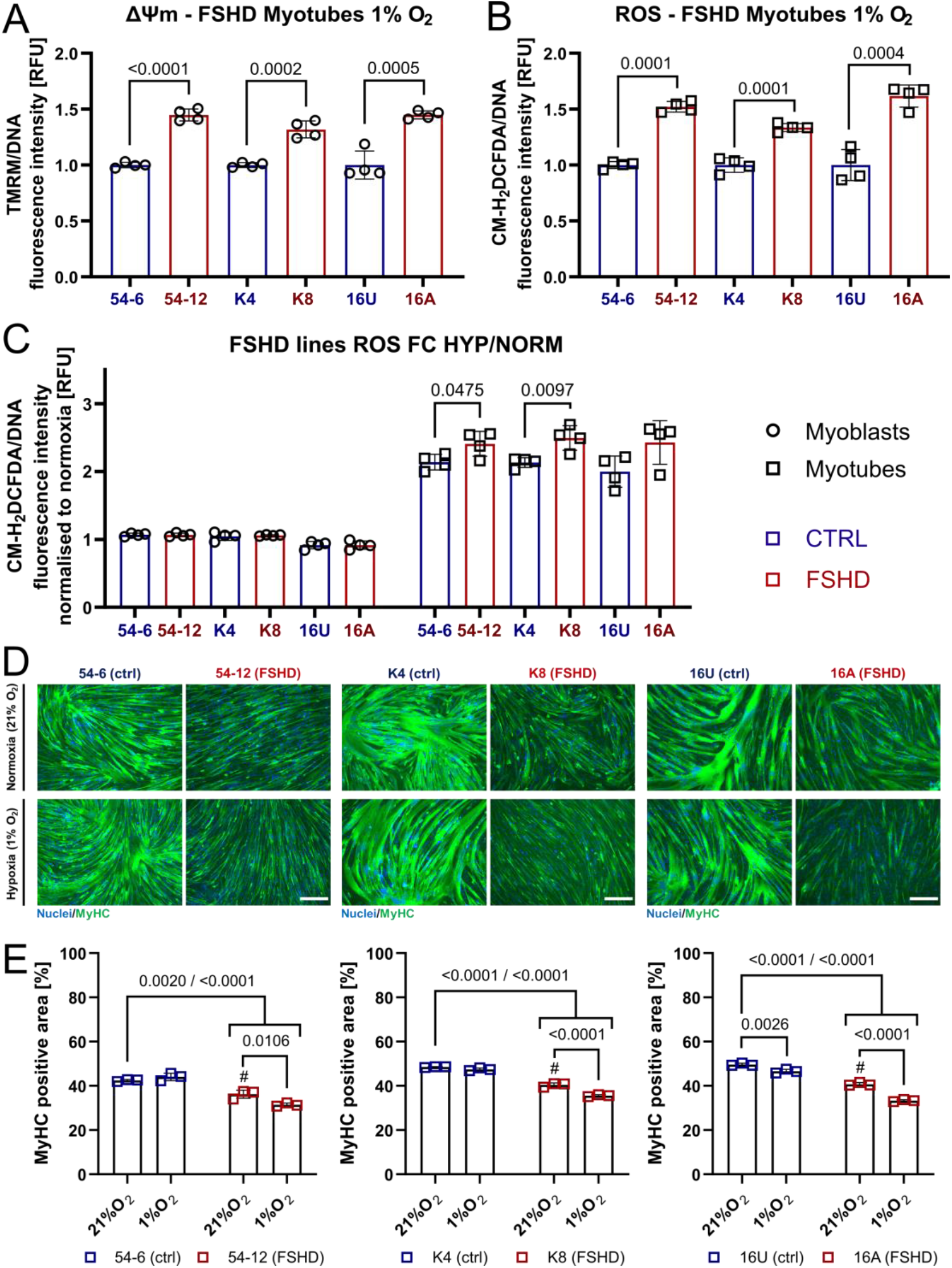
FSHD myogenesis is uniquely susceptible to hypoxia-induced oxidative stress. **(A, B)** FSHD patient myotubes (54-6 ctrl/54-12 FSHD; K4 ctrl/K8 FSHD; 16U ctrl/16A FSHD) maintain significantly elevated ΔΨm and ROS levels in hypoxia compared to isogenic/sibling control, but the difference is more pronounced than in normoxia (see Fig. 2A, B). **(C)** Hypoxia increases ROS levels in FSHD patient myotubes disproportionally compared to controls, which is not observed in FSHD myoblasts. **(D)** FSHD myotubes differentiated in hypoxia fail to properly metabolically adapt to low O_2_ availability, resulting in an aggravated hypotrophic phenotype. Hypoxic control myotubes are not affected (scale bar represents 250 μm). **(E)** Quantitation of MyHC-positive area as readout for myotube hypotrophy from immunofluorescence micrographs [# denotes statistical significance between hypoxic (1% O_2_) control myotubes and respective matched normoxic (21% O_2_) FSHD myotubes]. Data is mean ± s.d. from 3-4 wells each from a representative

### Mitochondria-targeted antioxidants efficiently rescue hallmarks of FSHD

Conventional, non-targeted antioxidant treatment can rescue aspects of FSHD pathology *in vitro* [44, 53, 71] and *in vivo* [55, 56], although with only moderate therapeutic efficiency. In addition, antioxidant rescue studies *in vitro* have only been performed under ambient O_2_ levels, so potential therapeutic effects on metabolic switching under low O_2_ availability are unknown. Having identified that mitoROS production by dysfunctional mitochondria upstream of oxidative stress disturbs FSHD myogenesis, specifically in hypoxia, we tested whether a more targeted antioxidant approach could more effectively rescue FSHD pathological hallmarks in hypoxic myotubes.

We selected 3 well-established antioxidant compounds with different modes of action and subcellular localisation: Vitamin C (VitC), a classic non-targeted antioxidant which is mostly retained in the cytoplasm; Coenzyme Q10 (CoQ10; ubiquinone-10), a lipophilic compound naturally involved in the mitochondrial ETC that can thus localize, to some extent, to mitochondria as well as other membranes; and mitoTEMPO, a mitochondria-targeted SOD-mimetic that accumulates several hundred-fold in mitochondria compared to cytoplasm due to its lipophilic cationic triphenylphosphonium moiety.

Initially, we tested for their ability to reduce ROS levels and ΔΨm after DUX4 induction in hypoxic iDUX4 myotubes, which produced the strongest morphological and redox phenotypes. VitC, CoQ10 and mitoTEMPO all reduced general ROS levels, with VitC and mitoTEMPO showing highest efficiency and almost identical reduction (Fig. 9A). This further emphasizes that mitoROS could be the primary trigger of oxidative stress, as ROS can leave the mitochondria in the form of H_2_O_2_ after dismutation by SOD, which would enable subsequent detoxification by cytoplasmic antioxidants like VitC. Notably though, only mitoTEMPO could reduce ΔΨm in response to DUX4 expression, demonstrating that (i) mitochondrial membrane polarisation is at least in part mediated by redox changes elicited by mitoROS and (ii) mitochondria-targeted antioxidant treatment may directly alleviate DUX4-induced mitochondrial dysfunction. Since mitoROS are a prerequisite for hypoxia signalling activation, only antioxidants capable of entering the mitochondria (mitoTEMPO and, to a much lesser extent, CoQ10) reduced iDUX4 myotube hypoxia (Fig. 9B). Interestingly, only mitoTEMPO was able to improve metabolic activity in DUX4 expressing myotubes, with VitC showing a detrimental effect, possibly due to pro-oxidant mechanisms. Importantly, all three compounds were able to rescue DUX4-induced myotube hypotrophy (Fig. 9C).

**Fig. 9:**
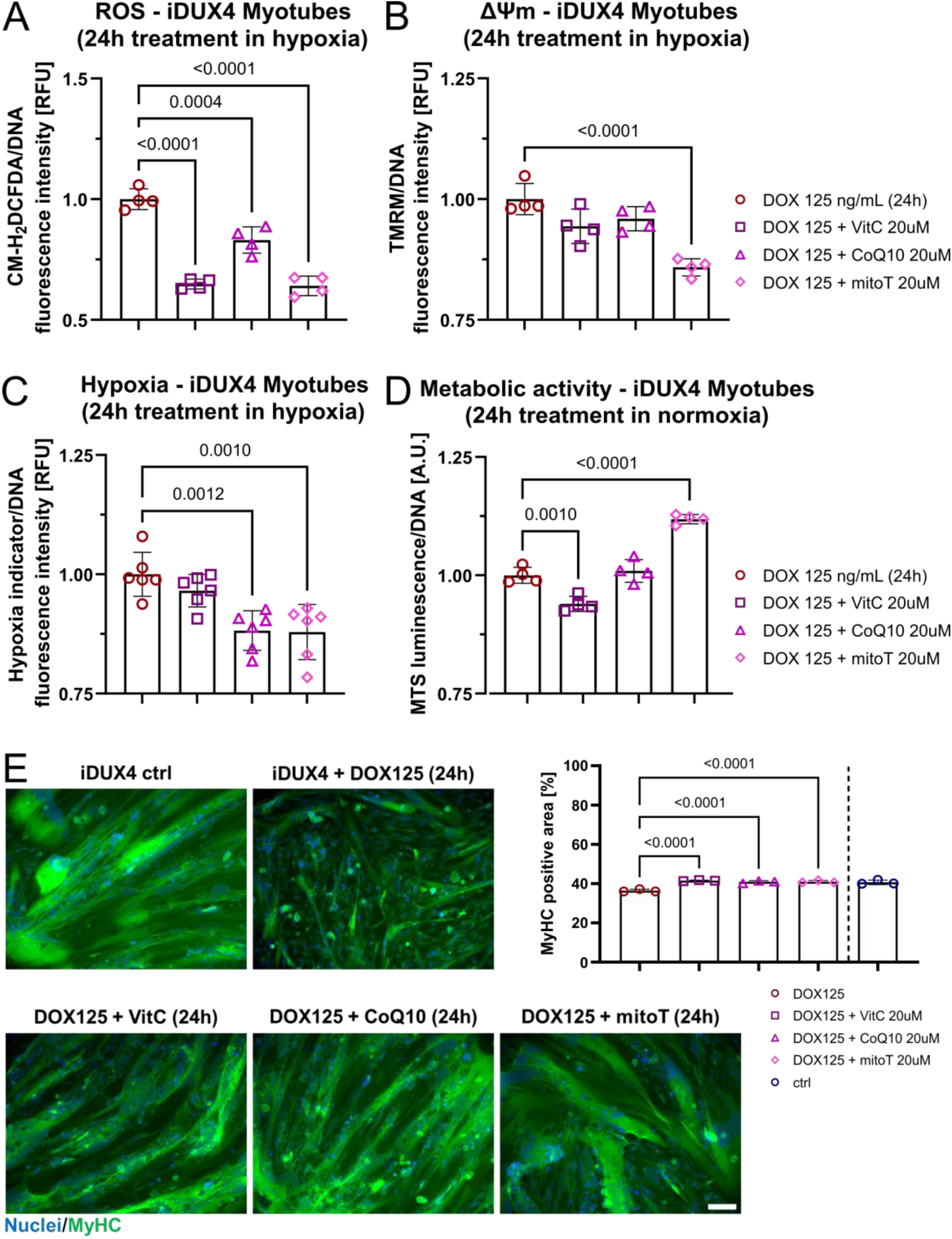
Mitochondria-targeted antioxidants more efficiently rescue DUX4-induced metabolic/hypoxic stress than conventional antioxidants. **(A, B)** Treatment of hypoxic iDUX4 myotubes with mitochondria-targeted (mitoTempo) or conventional (CoQ10, VitC) antioxidants effectively reduces ROS levels in response to DUX4 expression, but only mitoTempo normalises ΔΨm, reduces hypoxia and restores metabolic activity. **(C)** mitoTempo treatment phenotypically rescues DUX4 expressing iDUX4 myotubes in hypoxia with similar efficiency as non-targeted CoQ10 and VitC, emphasising central involvement of mitoROS as source of metabolic/hypoxic stress. Representative immunofluorescence micrographs are shown (scale bar represents 100 μm), as is quantitation of the MyHC positive area for each treatment group. Data is mean ± s.d. from 3-6 wells each from a representative experiment with *p* values as indicated.

We also tested these three antioxidants in patient-derived FSHD myotubes in hypoxia. All compounds reduced ROS levels with similar efficiency, but only CoQ10 and mitoTEMPO reduced ΔΨm, with mitoTEMPO the only compound capable of reducing FSHD myotube hypoxia (Fig. 10A). Notably, all three antioxidant compounds rescued FSHD myotube hypotrophy in hypoxia (Fig. 10B, C), demonstrating that targeting of redox-sensitive FSHD pathomechanisms has direct and beneficial effects on FSHD myogenesis under increased oxidative stress.

**Fig. 10:**
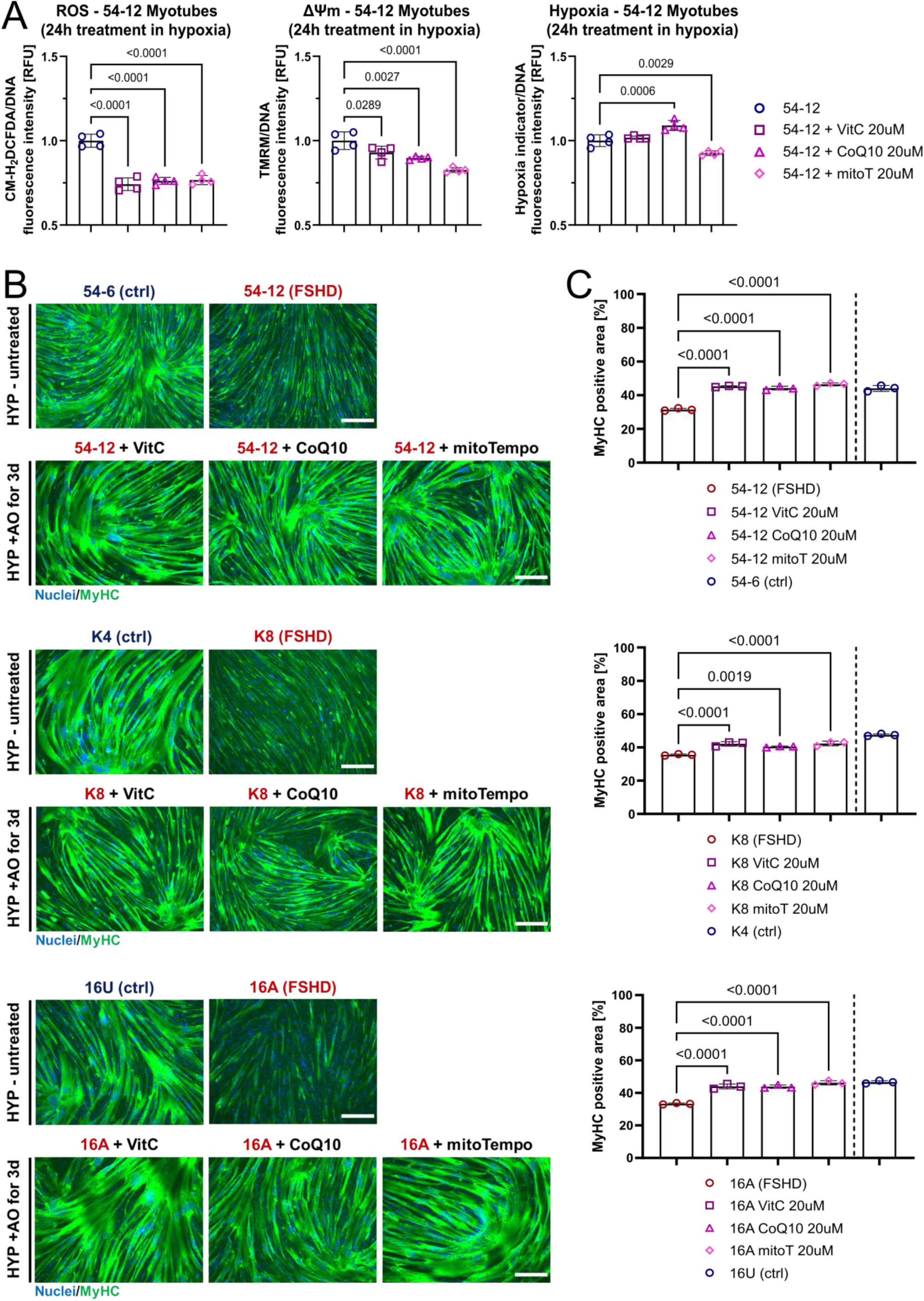
Mitochondria-targeted antioxidants alleviate oxidative stress and rescue aggravated FSHD myotube hypotrophy in hypoxia. **(A)** mitoTempo, CoQ10 and VitC demonstrate comparable efficiency in reducing ROS levels in hypoxic FSHD (54-12) patient myotubes, but mitoTempo shows highest ability to normalise ΔΨm and reduce hypoxia. **(B)** mitoTempo treatment phenotypically rescues FSHD myotube hypotrophy in hypoxia in 3 independent patient lines with similar efficiency as non-targeted CoQ10 and VitC. Representative immunofluorescence micrographs are shown [images of hypoxic controls from Fig. 8, as part of this experiment (scale bar represents 250 μm)]. **(C)** Quantitation of the MyHC positive area for each antioxidant treatment group compared to untreated FSHD myotubes (*K8 CoQ10 data from separate experiment, with given *p* value to that control). Untreated isogenic/sibling control included for comparison. Data is mean ± s.d. from 3-4 wells each from a representative experiment with *p* values as indicated.

Our analysis demonstrates that the efficiency of a given antioxidant compound in rescuing redox-sensitive FSHD pathomechanisms correlates with its ability to localise to mitochondria.

## DISCUSSION

This work for the first time provides a pathomechanistic model connecting two previously identified but under-studied central pathologic hallmarks of FSHD: mitochondrial dysfunction and disturbed hypoxia signalling. We identify that complex I-linked mitochondrial respiration is strongly and uniquely impaired in FSHD patient-derived muscle biopsies at the transcriptomic level. This is likely through mechanisms involving DUX4-induced redox perturbations triggering mitochondrial oxidative modifications and damage, and through more widespread transcriptional changes elicited by DUX4 in the nuclear genome.

Supraphysiological DUX4 levels in DUX4-inducible muscle cell models might lead to overestimation of DUX4 toxicity, as even low to moderate DOX concentrations already yield DUX4 levels much higher than found in FSHD patient-derived cell models and biopsies, where DUX4 is notoriously difficult to even detect [72]. We thus investigated redox changes in patient-derived models of FSHD as well as the LHCN-iDUX model, and found that changes in cellular RONS metabolism are remarkably consistent between both models, highlighting the central involvement of DUX4 in metabolic stress in FSHD. Specifically, FSHD mitochondria are characterised by hyperpolarised membranes, evident as a steady-state increase of ΔΨm in FSHD myoblasts/-tubes. Similarly, DUX4 expression elevates ΔΨm in a dose-dependent manner. Since the relationship between ΔΨm and mitoROS production is exponential [73], a small increase in ΔΨm can trigger high O_2_^·-^ production through drastically increased electron leakage from the respiratory chain. The notion that low level DUX4-induced changes in ΔΨm precedes elevated ROS levels and subsequent apoptosis (which is marked by a gradual depolarisation of the mitochondrial membrane) strongly suggest that the majority of ROS in FSHD stem from the respiratory chain, placing mitochondrial dysfunction upstream of oxidative damage.

While increased ROS levels in FSHD cells have been reported, we also found that elevated ΔΨm and ROS levels correlate inversely with NO· bioavailability. NO· and O_2_^·-^ can rapidly react to form the highly reactive oxidant ONOO^-^, suggesting that oxidative stress in FSHD also impairs the nitrosative system. DUX4 elicits mitochondrial lipid peroxidation, emphasizing the role of mitoROS-mediated NO· reduction and subsequent ONOO^-^ generation as source of oxidative mitochondrial insult, in line with a recent report [57]. Intriguingly, while DUX4 induction at low levels is marked by reduction in cellular NO·, ROS levels are initially unchanged but mitochondrial lipid peroxidation is already evident. Thus, NO· bioavailability through (mito)ROS presents an early event in the gradual disturbance of cellular redox control through DUX4. The rate constant for the reaction between NO· and O_2_·-is about one order of magnitude higher than that of O_2_^·-^ dismutation by SOD [74], so ONOO^-^ formation is an inevitable by-product of the mitochondrial respiratory chain. Since elevated mitoROS levels correlated with mitochondrial lipid peroxidation in response to DUX4 expression, DUX4-induced mitochondrial dysfunction likely confers metabolic stress through also acting indirectly on NO· signalling and bioavailability. Kinetics and compartmentalisation of interplay between O_2_·- and NO· are critical in transforming physiological regulatory effects of NO· to cytotoxic mechanisms through oxidative damage. Of note, as ONOO^-^ is a major mediator of oxidative stress through global changes in protein oxidation/nitration and lipid peroxidation [74].

Lipid peroxidation has been found in FSHD patient muscles [42], and there are membrane repair deficits in FSHD myoblasts and DUX4 expressing murine myofibres *ex vivo*, which are alleviated through antioxidant treatment [71]. Membrane lipid peroxidation is a comparably specific oxidative mechanism and it is unclear how more general oxidative/nitrosative protein modifications affect muscle function in FSHD. Oxidative and nitrosative stresses are potent activators of endoplasmic reticulum (ER)-stress and the unfolded protein response (UPR), which are increasingly recognized as important modulators of muscle health and disease [75]. *In vitro*, FSHD patient-derived myotubes are characterised by altered proteostasis, accompanied by aggregation of the transcriptional repressor TAR DNA-binding protein 43 (TDP-43) [76], a mechanism recently found to be mediated by TDP-43 S-nitrosylation [77]. Likewise, proteomics has identified a putative role of DUX4-induced ER-stress in FSHD pathogenesis, possibly through the well-characterised Heat Shock Protein Family A Member 5 (HSPA5)-Eukaryotic Translation Initiation Factor 2A (elF2a) axis [78]. Interestingly, this study also found changes in protein levels impacting on Golgi function and exocytosis, the latter regulated in part by primary and secondary RNS [79, 80]. While these studies suggest a broader metabolic stress (that may not be exclusively oxidative) at the core of FSHD pathogenesis, the role of the nitrosative system in FSHD pathogenesis is overlooked. In this respect, the versatile role of NO· in redox-regulation of skeletal muscle function through direct modulation of mitochondrial respiration and signalling [81–83], hypoxia response [84, 85] and apoptosis [86, 87] prompts research into the pathomechanistic contribution of perturbations of the nitrosative system in FSHD.

Having identified mitochondrial membrane polarisation and mitoROS at the core of DUX4-induced redox perturbance, our respirometric analysis revealed that mitochondrial dysfunction in response to DUX4 expression is uniquely conferred through complex I, a major source of mitoROS production [88]. Although several other sites of O_2_^·-^ production have been identified in mitochondria, complex I, alongside complex III, is considered the main driver of ROS production from the ETC [89], specifically at high pmf/ΔΨm. Although it is unclear to what extent individual ROS generating systems in the mitochondria contribute to oxidative stress in FSHD, as complex II-linked respiration was largely unaffected by DUX4, this suggests complex I as major source. Furthermore, as far as the substrate entry points into the ETC are concerned, complex I contributes to generation of ΔΨm while complex II does not under normal circumstances, linking mitochondrial membrane hyperpolarisation with altered complex I function. Chronic complex I inhibition causes mitochondrial membrane hyperpolarisation [90], an early event in mitoROS-mediated mitochondrial apoptosis and aging [91–93]. However, O_2_^·-^ production by complex I can also become excessively high during reverse electron transport (RET) from complex II during succinate-driven respiration [94], or when reversal of F_0_F_1_-ATPase is utilised to generate and restore ΔΨm through cytoplasmic ATP hydrolysis [95]. Our transcriptomic analysis from patient biopsies strongly supports complex I as the main trigger of mitochondrial dysfunction, identifying mitochondrial complex I assembly as the top enriched GOBP, along with strong enrichment for genes involved in mitochondrial gene expression, energy metabolism and, more general, respiratory chain complex assembly. Notably, we also identified robust transcriptional downregulation of all 13 protein-coding genes in the mitochondrial genome (of which 6 encode complex I subunits), indicating that mitochondrial gene expression is severely challenged by DUX4. To determine is whether this is a consequence of DUX4-induced oxidative mtDNA damage and thus gradual mtDNA instability, perturbance of the coordination between nuclear and mitochondrial gene expression during mitochondrial remodelling and biogenesis, or general reduction of mtDNA content. Interestingly, our GOBP analysis also revealed a strong link between pathways of mitochondrial (dys)function and nitrogen metabolism, with a considerable number of genes shared across both GO terms. Of note, NO· also participates in redox signalling pathways upstream of PGC1α-mediated mitochondrial biogenesis and turnover, which we have previously found dynamically repressed in FSHD myogenesis [48].

We observed a direct correlation between respiration and cellular hypoxia. DUX4 expression in myoblasts elicited a state resembling metabolic hypoxia [96], where decreased complex I-linked OXPHOS correlated with a decrease in hypoxia signalling through HIF1α. In myotubes, DUX4 had the opposite effect, with higher O_2_ consumption through increased complex I-linked OXPHOS further enhancing cellular hypoxia and HIF1α nuclear stabilisation. This correlation suggests that redistribution of O_2_ from the respiratory chain towards other O_2_-sensitive circuits (such as HIF1α activation) is a primary mechanism by which mitochondrial dysfunction affects hypoxia signalling in FSHD. Observed differences in OXPHOS between myoblasts and myotubes emphasise the importance of looking at DUX4-induced metabolic and redox changes over various developmental stages of myogenesis, since myoblasts mainly rely on glycolytic metabolism while terminally differentiated myotubes utilise OXPHOS as main energy source. This is highlighted by significant increase in mitochondrial mass and enzyme activity shortly after the onset of myogenic differentiation, and mitochondria have been identified as potent regulators of developmental and regenerative myogenesis [97]. However, the gradual switch from glycolytic to oxidative metabolism also renders myotubes a redox biological system quite different from myoblasts, as myotubes are much better equipped to deal with oxidative stress as a natural and inevitable byproduct of the bioenergetically more efficient OXPHOS. This difference may explain the increased susceptibility of FSHD myoblasts to oxidative stress compared to FSHD myotubes [36]. In this respect, the DOX concentrations used for respirometric analysis likely triggered much higher levels of oxidative stress/damage in iDUX4 myoblasts than in myotubes. Much higher concentrations of DOX were needed to trigger significantly elevated ROS levels in myotubes compared to myoblasts, and DOX treatment of myotubes with 62.5 ng/mL for 24h only yielded around 5% DUX4-postive myonuclei (as opposed to around 45% DUX-positive nuclei in myoblasts), which did not elicit a morphological myotube phenotype in normoxia. Nevertheless, pro-apoptotic iDUX4 myoblasts after 16h DOX treatment were marked by reduced complex I-linked OXPHOS and LEAK respiration, indicative of ONOO^-^-mediated impairment of mitochondrial membrane integrity, of complex I itself through irreversible S-nitrosylation, and, possibly, of mitochondrial transport proteins. Whether DUX4 initially affects ΔΨm and OXPHOS through perturbance of the mitochondrial transmembrane systems shuttling ions and metabolites, or directly through generation of mitochondrial oxidative stress, which then causes mitochondrial dysfunction, remains to be elucidated. A more thorough, kinetic respirometric analysis of DUX4-induced effects on OXPHOS will be needed to identify early versus late effects on mitochondrial (dys)function, specifically in myotubes where increased complex I-linked respiration could be an early event before the oxidative stress response is overwhelmed and complex I becomes irreversibly inhibited, as observed in the more oxidative stress-sensitive myoblasts.

Given that changes in OXPHOS directly affect cellular O_2_ consumption and are thus inevitably linked to hypoxia signalling activity, any impairment of mitochondrial function will impinge on the ability of cells to metabolically adapt to environmental hypoxia. Even though the relationship between hypoxia and mitoROS generation and its physiological significance remains controversial, there is accumulating evidence that myogenic cells produce higher amounts of ROS from the respiratory chain under hypoxic conditions [70, 98, 99]. We found that hypoxic culture increased ROS levels in non-induced iDUX4 myoblasts/myotubes, which were further increased by DUX4. Likewise, lower concentrations of DOX were sufficient to impair myotube formation in hypoxia compared to normoxia.

Since hypoxia did not overtly affect control myogenesis in non-induced cells, we hypothesise that DUX4 further increases hypoxia-induced oxidative stress through interference with hypoxic metabolic adaptation via HIF1α. mitoROS are a prerequisite for HIF1α activation [100, 101], and we found significantly elevated mitoROS levels in DOX treatment groups that produced a hypotrophic myotube phenotype. It is thus likely that, apart from O_2_ redistribution (through increased complex I-linked OXPHOS), DUX4 also causes enhanced mitoROS production from complex I to further aggravate myotube hypoxia by interference with metabolic adaptation through over-stimulation of HIF1α. Although the relation between hypoxia and ΔΨm is not completely understood, further elevation of ΔΨm by DUX4 in hypoxia (compared to normoxia) is probably the cause for enhanced oxidative stress in hypoxic myotubes through mitoROS. Similar to the DUX4 expression model, FSHD patient-derived myotubes maintain higher ΔΨm and ROS levels in hypoxia, the latter increasing disproportionally in hypoxic FSHD myotubes compared to normoxia. Hence, FSHD myotubes displayed aggravated hypotrophy when differentiated in hypoxia. Notably, this was not observed in FSHD myoblasts, where FSHD clones exhibit higher ROS levels in hypoxia (data not shown) and normoxia, but hypoxia did not increase ROS levels differentially.

Consistency between the results obtained in hypoxic DUX4 expressing and FSHD myotubes strongly suggests central involvement of DUX4 as trigger of oxidative stress and hypoxia sensitivity. Involvement of DUX4-induced oxidative stress as a negative regulator of FSHD myogenesis has mainly been shown in DUX4 overexpression systems, where antioxidants can alleviate myopathic phenotypes [28, 44, 71], but rarely in FSHD patient-derived cellular models, marked by much lower and sporadic DUX4 expression. In addition, conventional, non-targeted antioxidants have been investigated in all but one study [57]. To pinpoint involvement of mitoROS and hypoxia in FSHD pathogenesis, we used antioxidant compounds with different abilities to localise to mitochondria. VitC, CoQ10, mitoTEMPO all reduced ROS levels in DUX4 expressing hypoxic myotubes and rescued the hypotrophic myotube phenotype with comparable efficiency, but only mitochondria-targeted mitoTEMPO reduced ΔΨm and hypoxia, while restoring metabolic activity. These findings suggest that (i) the respiratory chain is a major contributor to oxidative stress through mitoROS (specifically when metabolic switching is required in response to hypoxia), (ii) mitoROS initially released into the mitochondrial matrix or intermembrane space eventually escape into the cytoplasm as H_2_O_2_ to further interfere with the cellular redox balance, and (iii) mitoROS are involved in over-stimulation of HIF1α in DUX4 expressing myotubes. Likewise, antioxidant treatment rescued the hypotrophic phenotype of hypoxic FSHD myotubes in three separate patient-derived myoblast lines, where FSHD myotubes had shown increased sensitivity to hypoxia. Notably, mitoTEMPO was most efficient in normalising ΔΨm and reducing hypoxia through its ability to enrich within mitochondria, yet no difference in the ability of the antioxidants to rescue FSHD myogenesis was observed, further underscoring that oxidative stress in FSHD stems predominantly from mitoROS.

In summary, we have identified mitochondrial dysfunction followed by elevated generation of mitoROS as a primary mechanism by which DUX4 causes oxidative stress in FSHD (Fig. 11). We provide a link between the redox biological changes elicited through disturbed RONS metabolism and hypoxia sensitivity, and show complex I dysfunction with enhanced mitoROS production from the respiratory chain as a main trigger of DUX4-mediated pathogenesis. We further demonstrate that mitochondria-targeted antioxidants are more effective in alleviating aspects of disturbed myotube metabolism compared to conventional antioxidants, highlighting involvement of the respiratory chain as a source of ROS. Given the moderate clinical outcomes in FSHD patients seen in clinical trials employing non-targeted antioxidants [56, 102], mitochondria-targeted compounds should be a more efficient approach, as they target ROS directly at the site of generation and require much lower concentrations to be effective *in vivo* while interfering less with physiologically important redox pathways [103]. Indeed, mitochondria-targeted antioxidants have been proven safe in humans [104], with beneficial effects in phase II clinical trials for hepatitis C [105] and dry eye treatment [106].

**Fig. 11:**
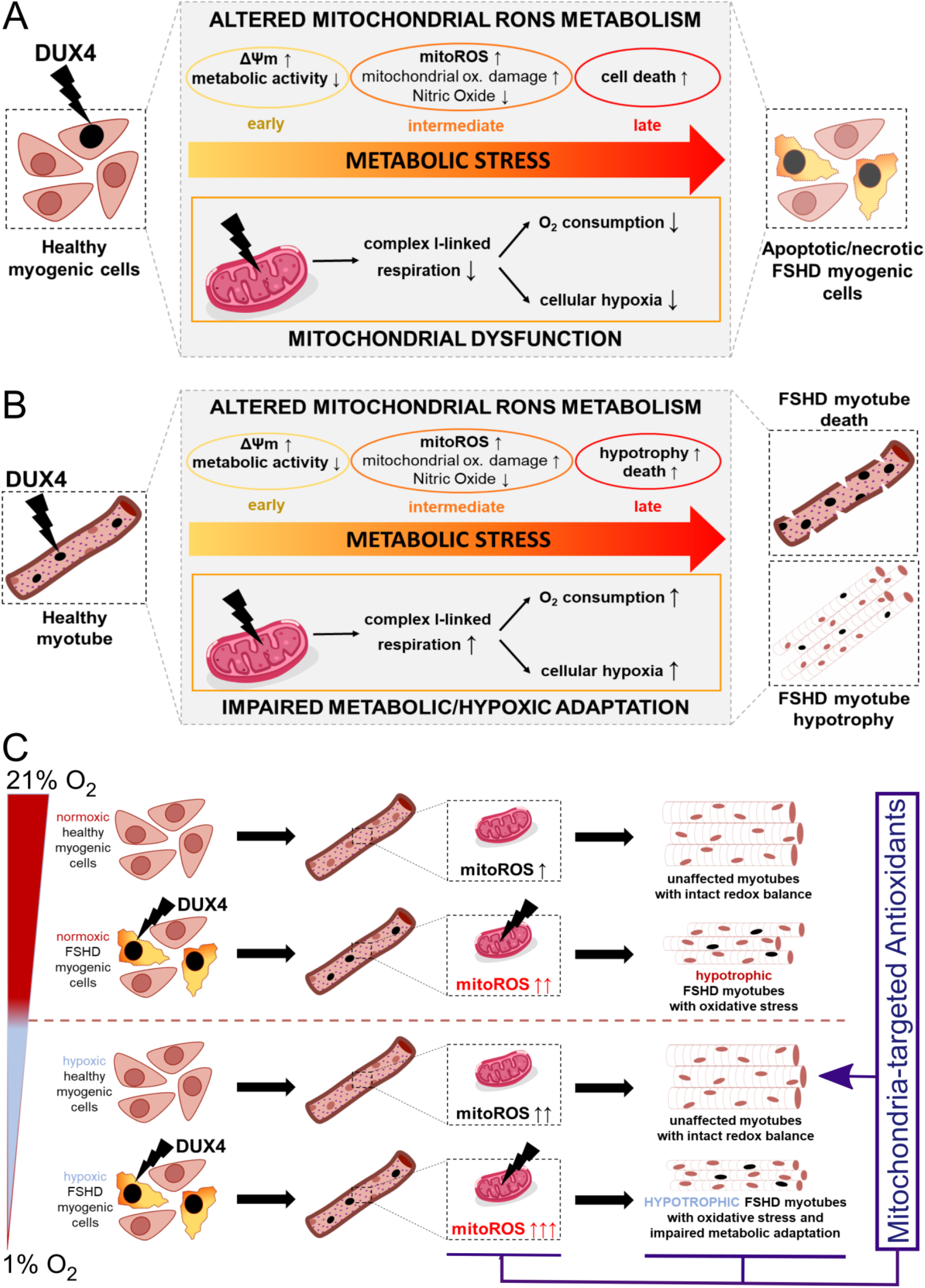
Mechanisms of metabolic stress generation in FSHD. **(A)** DUX4 triggers metabolic stress in myoblasts through alterations in mitochondrial RONS metabolism and function. Hyperpolarisation of the mitochondrial membrane is an early event in response to DUX4, followed by mitochondrial oxidative damage through enhanced mitoROS formation from the respiratory chain. Mitochondrial dysfunction is conferred through reduced complex I-linked respiration, affecting hypoxia signalling through redistribution of O_2_. **(B)** DUX4 also triggers oxidative damage and altered mitochondrial RONS metabolism driven by high ΔΨm in myotubes, resulting in myotube hypotrophy and apoptosis. Increased O_2_ consumption via complex I and enhanced mitoROS formation both trigger hypoxia in myotubes. **(C)** DUX4-induced redox changes challenge mitochondrial health and function through altered RONS metabolism, and thus interfere with metabolic adaptation to environmental hypoxia. Enhanced mitoROS formation triggered by DUX4 in normoxic myotubes is further increased in hypoxia, concomitant with aggravation of the hypotrophic myotube phenotype. Notably, FSHD myotubes are uniquely sensitive to hypoxia-induced metabolic/oxidative stress, whereas control myotubes adapt their metabolism to prevent oxidative stress through excess mitoROS. Given that mitoROS-induced oxidative stress is a main driver of myotube hypotrophy in hypoxia, mitochondrial-targeted antioxidants alleviate FSHD phenotypes more efficiently than conventional non-targeted antioxidants. Thus, affected mitochondria in FSHD are a primary trigger of muscle loss associated with the disease, specifically when metabolic adaptation to varying O_2_ availability is required.

Mitochondria-targeted antioxidants will also be useful to elucidate the mechanistic role of disturbed mitochondrial ROS metabolism in FSHD pathogenesis. Of significance, two prominent extra-muscular features of FSHD, retinal telangiectasia and sensorineural hearing loss, have also been linked with oxidative stress and mitochondrial dysfunction [107, 108]. Both retina and cochlea are characterized by high metabolic activity, and, specifically, the central involvement of OXPHOS deficits in the pathophysiology of hearing loss is well characterized [109]. While the beneficial effects of conventional antioxidants in treatment of hearing loss remain controversial [110], mitochondria-targeted antioxidants have yielded promising results in animal studies [111].

It is crucial to now identify how DUX4 challenges cellular redox pathways to understand the extent of the metabolic stresses in FSHD. Deeper understanding of OXPHOS-related pathomechanisms will not only inform novel therapeutics such as mitochondria-targeted antioxidants, Szeto-Schiller (SS) peptides or mild uncouplers [112, 113], but will likely also reveal novel aspects of FSHD aetiology. DUX4 affects more than 200 genes “indirectly” by oxidative stress in normoxic myoblasts [57]. Given discrepancies between the available transcriptomic and proteomic data sets [78], investigating DUX4-induced transcriptional changes will not allow conclusions regarding the FSHD metabolome. Dynamic transcriptomic-metabolomic analyses of FSHD myogenesis under O_2_ tensions to model physioxia and hypoxia are needed to find novel pathomechanisms related to metabolic adaptation undetectable in normoxia. This will not only decipher mechanisms of metabolic stress related to hypoxia, but will also identify whether the redox-sensitive core oxidative metabolic pathways providing substrates for OXPHOS are uniquely affected, in addition to the mitochondrial respiratory chain. Since FSHD patients show signs of both muscular and systemic oxidative stress/damage, evaluation of mechanisms described here in non-myogenic FSHD models will be useful to investigate how aspects of DUX4-induced redox perturbance affects cell and tissue function in these models.

## MATERIALS AND METHODS

### Transcriptomic analyses

For differential expression (DE) analysis, RNA-Seq data (gene counts) of muscle biopsies from 6 severely affected FSHD patients and 9 unaffected individuals were obtained from GSE115650 [58]. Data processing was performed as recently described [114]. Briefly, filtering for lowly expressed genes (CPM < 1) and further biotype filtering was performed in R with the Bioconductor packages NOISeq [115] and biomaRt [116] to remove highly expressed mitochondrial and ribosomal RNA. DE analysis was performed using the Bioconductor package edgeR [117]. Specifically, a negative binomial generalized log-linear model (glmfit) was fitted to the read counts and the likelihood ratio test (glmLRT) was conducted for each comparison of interest. The Benjamini-Hochberg false discovery rate (FDR) cut-off was set at 0.05. The versions of all relevant Bioconductor packages were compatible with R v3.5.3.

Heatmap depicting expression levels of the 887 unique genes listed in all GOs referring to superclusters Mitochondrial Activity & Organisation, Response to oxidative stress and oxygen levels and metabolism of nitrogen compound, was generated using Morpheus (https://software.broadinstitute.org/morpheus/) applying the function “One plus Log2” followed by hierarchical clustering.

For pathway and gene set analysis, we used WebGStalt (www.webgestalt.org) to assess gene ontology terms that showed significant enrichment in the list of DEG (Overrepresentation Enrichment Analysis). The enrichment for each GO Biological Process (GOBP) term was considered statistically significant if the adjusted p-value (FDR) was lower than 0.05. Only the top 30 GOs are reported in figure 1. Cytoscape v. 3.7.2 [118] was used to visualise relevant biological networks of enriched GOBPs, together with EnrichmentMap and AutoAnnotate applications. Several layout parameters were tuned to achieve the current Cytoscape visualization.

### Cell culture and myogenic differentiation

The three immortalised FSHD patient-derived cellular models were the isogenic ‘54’ series derived from the biceps of a male mosaic FSHD1 patient [119], where 54-6 (13 D4Z4 repeats) is the uncontracted control clone and 54-12 (3 D4Z4 repeats) the contracted FSHD clone; the isogenic ‘K’ (K271TA44) series from the tibialis anterior of a mosaic FSHD1 patient, with K4 the uncontracted control and K8 the contracted FSHD clone; and the ‘16’ series, a sibling-matched immortalised model derived from biceps muscle [120], where 16A is the contracted FSHD line and 16U the uncontracted control line from a first-degree relative. The DUX4-inducible myoblast line (iDUX4) was LHCN-M2-iDUX, on the LHCN-M2 myoblast background [121]. DUX4 expression was induced by doxycycline (DOX; Sigma Aldrich, Dorset, UK).

Human myoblast lines were cultured in Skeletal Muscle Cell Growth Medium (Promocell, Heidelberg, Germany) supplemented with 20% foetal bovine serum (FBS; ThermoFisher Scientific, MA, USA), 50 μg/ml fetuin (bovine), 10 ng/ml epidermal growth factor (recombinant human), 1 ng/ml basic fibroblast growth factor (recombinant human), 10 μg/ml insulin (recombinant human), 0.4 μg/ml dexamethasone (all supplemented as PromoCell SupplementMix) and 50 μg/ml gentamycin (Sigma Aldrich) in a humidified incubator at 37°C with 5% CO_2_. Myoblast lines were kept subconfluent in routine culture, and passaged at maximum 70% confluency.

To induce differentiation, myoblasts were washed twice with phosphate buffered saline (PBS) and placed in Dulbecco’s Modified Eagle Medium (DMEM) GlutaMax (ThermoFisher Scientific) supplemented with 0.5% FBS, 10 ug/mL recombinant human insulin (Sigma-Aldrich) and 50 ug/mL gentamycin. Where applicable, Vitamin C (L-ascorbic acid), Coenzyme Q10 (ubiquinone) and mitoTEMPO (all from Sigma Aldrich) were added to the differentiation medium at final concentrations of 20 uM for each antioxidant compound.

Cell culture under hypoxic conditions was performed in a commercially available humidified hypoxia chamber (STEMCELL technologies, Cambridge, UK) in an incubator at 37°C. To keep O_2_ tension constant, the hypoxia chamber was injected with a gas mixture of 1% O_2_, 5% CO_2_ and 94% N_2_ (Air Liquide, London, UK) at continuous flow rate of approximately 1L/min. For medium changes in hypoxic cultures, e.g. for RONS and hypoxia measurements or antioxidant treatment, all medium, buffer and reagent solutions were pre-conditioned in hypoxia for at least the experimental duration, or, for longer experiments in differentiation, for at least 24 h.

### RONS measurements

For RONS measurements in proliferating myoblasts, 10000 cells/well were seeded into black, clear-bottom polystyrene 96-well plates (Corning®, Sigma Aldrich) and assayed 24 h later. For measurements in hypoxia, 5000 cells/well were seeded, transferred into hypoxia the following day and assayed 24 h later, with reagents preconditioned in hypoxia for 24 h. To assess RONS levels in differentiated myotubes, 50000 cells/well were seeded into black, clear-bottom polystyrene 96-well plates, and switched to differentiation 24 h later. RONS measurements were performed in myotubes after 3 days of differentiation (in normoxia or hypoxia). RONS probes used and staining conditions are detailed in table 1.

**Table 1:** fluorescent probes used for RONS measurements.

| Species | Probe | Supplier | Conc. | Staining duration | $\lambda$ Ex/Em |
| --- | --- | --- | --- | --- | --- |
| General<br>(cytoplasmic)<br>ROS | CM-H<sub>2</sub>DCFDA | ThermoFisher<br>Scientific | 5 μM | 30 min (in HBSS) | 485/520 |
| NO· | DAF-FM<br>Diacetate | ThermoFisher<br>Scientific | 5 μM | 30 min (in HBSS),<br>followed by 20 min in<br>growth medium for de-esterification | 485/520 |
| mitoROS | MitoTracker<sup>®</sup> Red<br>CM-H<sub>2</sub>XROS | ThermoFisher<br>Scientific | 250 nM | 30 min (in HBSS) | 584/620 |

RONS measurements were performed in Hank’s Balanced Salt Solution (HBSS) supplemented with Ca^2+^ and Mg^2+^ (Sigma Aldrich). Cells were washed twice with HBSS, followed by incubation with the respective fluorescent RONS probe and HOECHST33342 (0.5 ug/mL; ThermoFisher Scientific) in HBSS for 30 min in the dark. After staining, cells were washed twice with HBSS (with the exception of DAF-FM DA, where a 20 min incubation step in growth medium for de-esterification of the probe was performed prior to washing) and fluorescence intensity was measured on a POLARStar Omega microplate reader (BMG Labtech, Aylesbury, UK) in orbital averaging scan mode (20 flashes per well, scan diameter 4 mm). For normalization to cell number, RONS probe fluorescence was normalized to DNA content as simultaneously assessed via HOECHST 33342 fluorescence in the same well.

### Assessment of ΔΨm and mitochondrial lipid peroxidation

ΔΨm was assessed by Tetramethylrhodamine methyl ester (TMRM; Sigma Aldrich) staining in black clear-bottom polystyrene 96-well plates. Briefly, cells were washed twice with HBSS and subsequently stained with 100 nM TMRM and 0.5 ug/mL HOECHST33342 in HBSS for 30 min at 37°C in the dark. After staining, cells were washed twice with HBSS and fluorescence intensity was measured on a POLARStar Omega microplate reader in orbital averaging scan mode (20 flashes per well, scan diameter 4mm). For normalization to cell number, TMRM fluorescence was normalized to DNA content as simultaneously assessed via HOECHST 33342 in the same well.

Mitochondrial lipid peroxidation was measured with the ratiomeric C11-BODIPY derivative MitoPerOx (Abcam, Cambridge, UK). For iDUX4 myoblasts in proliferation, 25000 cells/well were seeded into black clear-bottom polystyrene 96-well plates and, after 24h, induced to express DUX4 for 16h. For measurements in iDUX4 myotubes, 50000 cells/well were seeded and differentiation induced 24h later. After 36h, when myotube formation was evident, DUX4 expression was induced for 24h with subsequent assaying. For staining, cells were washed twice with HBSS and incubated with 100 nM MitoPerOx probe and 0.5 ug/mL HOECHST33342 in HBSS for 45 min at 37°C in the dark. After staining, cells were washed twice with HBSS and fluorescence intensity was measured on a POLARStar Omega microplate reader in orbital averaging scan mode (20 flashes per well, scan diameter 4mm). Mitochondrial lipid peroxidation was quantified by calculating the ratio between fluorescence intensity at em520/em590 after excitation at 488 nm, with subsequent normalization to DNA content as simultaneously assessed via HOECHST 33342.

### Metabolic activity measurement

Metabolic activity was measured using the luminescence RealTime-Glo^TM^ MT Cell Viability assay (Promega, Southampton, UK) with a modified protocol. In proliferation, 5000 iDUX4 cells/well were seeded into opaque walled polystyrene 96-well plates (Corning®, Sigma Aldrich). 24h later, DUX4 expression was induced and assay substrates were added simultaneously for longitudinal assaying for up to 16h according to the manufacturer’s protocol. Luminescence was read with a Mithras LB940 multimode microplate reader (Berthold Technologies, Bad Wildbad, Germany) at 0.1s exposure, and metabolic activity was calculated as the ratio between luminescent signal and DNA content, as assessed through HOECHST33342 fluorescence from sister cultures undergoing the same treatment. For iDUX4 myotubes, the assay was used in an end-point format. Briefly, 50000 cells/well were seeded into opaque 96-well plates and induced to differentiate 24h later. After 36h, DUX4 expression was induced for 24h (with simultaneous antioxidant supplementation where applicable) and, at the end of the experimental treatment duration, assay substrates were added according to the manufacturer’s protocol. Luminescence was read after 3h incubation with the MT assay substrates, and metabolic activity calculated as described above.

### Apoptosis and Necrosis assaying

Apoptosis and Necrosis were assayed with the combined luminescence/fluorescence RealTime-Glo^TM^ Annexin V Apoptosis and Necrosis assay (Promega), according to the manufacturer’s instructions. 5000 iDUX4 cells/well were seeded into black clear-bottom polystyrene 96-well plates. 24h later, cells were induced to express DUX4 and assay substrates were added simultaneously for longitudinal assaying of apoptosis/necrosis for up to 24h according to the manufacturer’s protocol. Apoptosis was quantified as luminescent signal intensity, as assessed on a Mithras LB940 multimode microplate reader (0.1s exposure), and necrosis was quantified as fluorescence intensity (ex488/em520), as assessed on a POLARStar Omega microplate reader in orbital averaging scan mode (20 flashes per well, scan diameter 4mm).

### High resolution respirometry (HRR)

Respirometric analysis was performed on an Oroboros O2k oxygraph (Oroboros Instruments, Innsbruck, Austria). In proliferation, 4 x 10^6^ viable iDUX4 myoblasts were used per experiment (after 16h of DUX4 expression). Cell viability was assessed by trypan Blue (Sigma Aldrich) staining prior to each respirometric analysis, and oxygen consumption rate normalised to cell number. For respirometric analysis of iDUX4 myotubes, 5.86 x 10^6^ cells were seeded per 75 cm^2^ cell culture flasks (Nunc^TM^, ThermoFisher Scientific), and switched to differentiation 24h later. After 36h, after myotubes had formed, DUX4 expression was induced for another 24h, and measurements performed from myotubes harvested by trypsination. To account for differential cell/myotube amounts, DNA quantitation of sister cultures undergoing the same treatment was performed by HOECHST33342 staining, as described above, and HEOCHST33342 fluorescence subsequently used as normaliser for oxygen consumption rate.

Respiratory states were analysed through substrate-uncoupler-inhibitor-titration (SUIT) as follows: to assess Complex I-and Complex II-linked respiration, cells were incubated in a buffer containing 80 mM KCl, 5 mM KH_2_PO_4_, 50 mM 3-(N-morpholino)propanesulfonic acid, 1 mM ethylene glycol-bis(2-aminoethylether)-N,N,N′,N′-tetraacetic acid, and 1 mg/mL fatty acid-free bovine serum albumin (pH 7.4, 37 °C) [122], and permeabilized with digitonin (8 µM). State 2 respiration (LEAK) was induced either by the addition of 5 mM glutamate and 5 mM malate (complex I), 10 mM pyruvate (complex I), or 10 mM succinate in the presence of 1.4 µM rotenone (complex II). State 3 respiration (OXPHOS) was stimulated by the addition of 1 mM adenosine diphosphate. Maximum electron transfer system capacity (max. ETS) was measured by titration of carbonyl cyanide-4-(trifluoromethoxy)phenylhydrazone in steps of 0.1 µM. To calculate respiratory control ratio (RCR), State 3 respiration (OXPHOS) was divided by State 2 respiration (LEAK). Oxygen consumption rates were obtained by calculating the negative time derivative of the measured oxygen concentration.

### Cellular hypoxia measurement

Cellular oxygenation was determined with the fluorescent O_2_-sensitive hypoxia indicator Image-IT^TM^ Green Hypoxia Reagent (ThermoFisher Scientific) according to the manufacturer’s instructions. Cells seeded in black clear-bottom polystyrene 96-well plates were incubated with 5 uM Green Hypoxia Reagent in growth or differentiation medium, respectively, for 30 min at 37°C in the dark. Subsequently, cells were washed with HBSS, treated with DOX (for DUX4 expression in iDUX4 cells) and/or antioxidants and transferred to hypoxia for the appropriate remaining culture period (e.g. 16h for iDUX4 myoblasts induced to express DUX4 or 24h for antioxidant treatment in iDUX4 or FSHD myotubes). To quantify cellular hypoxia, cells were washed twice with HBSS and hypoxia indicator fluorescence intensity (ex488/em520) measured on a POLARStar Omega microplate reader in orbital averaging scan mode (20 flashes per well, scan diameter 4mm). Afterwards, DNA content was determined for normalisation by staining with 0.5 ug/mL HOECHST33342 in HBSS for 30 min at 37°C in the dark, and subsequently reading HOECHST33342 fluorescence intensity. For visualisation of Green Hypoxia Reagent fluorescence, fluorescence micrographs were taken on a Zeiss Axiovert 200M epifluorescence microscope using a Zeiss AxioCam HRm and AxioVision 4.4 software (Zeiss, Jena, Germany).

### Immunofluorescence microscopy

For immunolabelling, cells were fixed with 4% paraformaldehyde/PBS (Alfa Aesar, Heysham, UK) for 10 min, washed three times with PBS for 5 min each, then permeabilised with 0.1% Triton-X/PBS (Sigma Aldrich) for 15 min and washed three times with PBS again. Blocking was performed in 5% normal goat serum (GS)/PBS (Dako, Glostrup, Denmark) for 60 min, and, after three washes with PBS, cells were incubated with primary antibody in 1% GS/PBS on a rocker overnight at 4°C. Cells were then washed three times with PBS, and incubated with secondary antibody in 1% GS/PBS for 60 min in the dark at room temperature. After three more washes with PBS, nuclei were stained with 0.5 ug/mL HOECHST33342 in PBS for 10 min, washed again with PBS and imaged on a Zeiss Axiovert 200M epifluorescence microscope using a Zeiss AxioCam HRm and AxioVision 4.4 software. Primary antibodies were mouse anti-MyHC (1:400; MF-20; Developmental Studies Hybridoma Bank, IA, USA), mouse anti-DUX4 (1:500; clone 9A12; Merck Millipore, Croxley Park, UK) and rabbit anti-HIF1α (1:500; EP1215Y, Abcam). Secondary antibodies were goat anti-mouse IgG (H+L) AlexaFluor-488 (1:500, A-11001, ThermoFisher Scientific) and goat anti-rabbit IgG (H+L) AlexaFluor-594 (1:500, A-11012, ThermoFisher Scientific).

### Determination of MyHC-positive area and differentiation index

Image analysis was performed from MyHC immunofluorescence micrographs with a custom-made high-throughput image analysis software, as recently described [48]. For determination of the MyHC-positive area the channel displaying MyHC was passed through a low pass filter and a size filter was applied to remove background labelling in the binarized image. The positive proportion of the image was then quantified as MyHC-positive area. The mean MyHC-positive area per well was calculated as the average from three representative images per well. Likewise, differentiation index was automatically determined from the same images as the percentage of nuclei within the MyHC-positive area.

### Statistical analysis

Statistical analysis was performed using GraphPad Prism version 9.1.2 for Windows (GraphPad Software, San Diego, CA, USA; www.graphpad.com). Experiments were performed at least 3 independent times, with detailed N numbers and technical replicates given in each figure. Variance between groups was compared using a Brown Forsythe test and revealed no significant difference. A comparison between two groups was performed using an unpaired homoscedastic two-tailed student’s *t*-test. A comparison of more than two groups was performed using a one-way analysis of variance (ANOVA) followed by either Dunnett’s post-test when different groups were compared with the control group or Tukey’s post-test when different groups were compared with each other. *p* < 0.05 was considered significantly different, with *p* values indicated for each significant comparison in the figures.

## Acknowledgements

We would like to thank: Vincent Mouly (Center for Research In Myology, UMRS 974 Sorbonne Université-INSERM, Paris, France) and Charles P. Emerson (Wellstone Muscular Dystrophy Program, University of Massachusetts Medical School, MA, USA) for providing immortalised myoblast lines; Alexandra Belayew, Luca Pinton, Terje Wimberger, Christian Oeffel for helpful discussion; Christopher R. S. Banerji and Sara Badodi for assistance in RNAseq analysis; Johannes Oesterreicher, Nadja Milivojev and Sergejs Zavadskis for useful input on RONS measurements.

## Author contributions

The research study was conceived and designed by PH, AVK and PSZ, with input from AT, AED, AK and JG. Experiments were performed, and data acquired/analysed, by PH, MG, AW, EE, JP and THN. Human myoblast lines were provided by KM. The manuscript was written by PH and PSZ, with support from other authors. The main funding for the work was secured by PSZ and PH.

## REFERENCES

1. Deenen, J.C., et al., Population-based incidence and prevalence of facioscapulohumeral dystrophy. Neurology, 2014. 83(12): p. 1056–9.

2. Sposito, R., et al., Facioscapulohumeral muscular dystrophy type 1A in northwestern Tuscany: a molecular genetics-based epidemiological and genotype-phenotype study. Genet Test, 2005. 9(1): p. 30–6.

3. Tawil, R. and S.M. Van Der Maarel, Facioscapulohumeral muscular dystrophy. Muscle Nerve, 2006. 34(1): p. 1–15.

4. Tawil, R., S.M. van der Maarel, and S.J. Tapscott, Facioscapulohumeral dystrophy: the path to consensus on pathophysiology. Skelet Muscle, 2014. 4: p. 12.

5. Pandya, S., W.M. King, and R. Tawil, Facioscapulohumeral dystrophy. Phys Ther, 2008. 88(1): p. 105–13.

6. Lutz, K.L., et al., Clinical and genetic features of hearing loss in facioscapulohumeral muscular dystrophy. Neurology, 2013. 81(16): p. 1374–7.

7. Fitzsimons, R.B., Retinal vascular disease and the pathogenesis of facioscapulohumeral muscular dystrophy. A signalling message from Wnt? Neuromuscul Disord, 2011. 21(4): p. 263–71.

8. Sakellariou, P., et al., Mutation spectrum and phenotypic manifestation in FSHD Greek patients. Neuromuscul Disord, 2012. 22(4): p. 339–49.

9. Tawil, R., et al., Extreme variability of expression in monozygotic twins with FSH muscular dystrophy. Neurology, 1993. 43(2): p. 345–8.

10. Tonini, M.M., et al., Asymptomatic carriers and gender differences in facioscapulohumeral muscular dystrophy (FSHD). Neuromuscul Disord, 2004. 14(1): p. 33–8.

11. Calandra, P., et al., Allele-specific DNA hypomethylation characterises FSHD1 and FSHD2. J Med Genet, 2016. 53(5): p. 348–55.

12. Gabriels, J., et al., Nucleotide sequence of the partially deleted D4Z4 locus in a patient with FSHD identifies a putative gene within each 3.3 kb element. Gene, 1999. 236(1): p. 25–32.

13. Hewitt, J.E., et al., Analysis of the tandem repeat locus D4Z4 associated with facioscapulohumeral muscular dystrophy. Hum Mol Genet, 1994. 3(8): p. 1287–95.

14. van Deutekom, J.C., et al., FSHD associated DNA rearrangements are due to deletions of integral copies of a 3.2 kb tandemly repeated unit. Hum Mol Genet, 1993. 2(12): p. 2037–42.

15. van Overveld, P.G., et al., Hypomethylation of D4Z4 in 4q-linked and non-4q-linked facioscapulohumeral muscular dystrophy. Nat Genet, 2003. 35(4): p. 315–7.

16. Wijmenga, C., et al., Chromosome 4q DNA rearrangements associated with facioscapulohumeral muscular dystrophy. Nat Genet, 1992. 2(1): p. 26–30.

17. Kowaljow, V., et al., The DUX4 gene at the FSHD1A locus encodes a pro-apoptotic protein. Neuromuscul Disord, 2007. 17(8): p. 611–23.

18. Lemmers, R.J., et al., A unifying genetic model for facioscapulohumeral muscular dystrophy. Science, 2010. 329(5999): p. 1650–3.

19. Tupler, R., et al., Monosomy of distal 4q does not cause facioscapulohumeral muscular dystrophy. J Med Genet, 1996. 33(5): p. 366–70.

20. Lemmers, R.J., et al., Digenic inheritance of an SMCHD1 mutation and an FSHD-permissive D4Z4 allele causes facioscapulohumeral muscular dystrophy type 2. Nat Genet, 2012. 44(12): p. 1370–4.

21. van den Boogaard, M.L., et al., Mutations in DNMT3B Modify Epigenetic Repression of the D4Z4 Repeat and the Penetrance of Facioscapulohumeral Dystrophy. Am J Hum Genet, 2016. 98(5): p. 1020–1029.

22. Hamanaka, K., et al., Homozygous nonsense variant in LRIF1 associated with facioscapulohumeral muscular dystrophy. Neurology, 2020. 94(23): p. e2441–e2447.

23. Jansz, N., et al., The Epigenetic Regulator SMCHD1 in Development and Disease. Trends Genet, 2017. 33(4): p. 233–243.

24. Wallace, L.M., et al., DUX4, a candidate gene for facioscapulohumeral muscular dystrophy, causes p53-dependent myopathy in vivo. Ann Neurol, 2011. 69(3): p. 540–52.

25. Xu, H., et al., Dux4 induces cell cycle arrest at G1 phase through upregulation of p21 expression. Biochem Biophys Res Commun, 2014. 446(1): p. 235–40.

26. Knopp, P., et al., DUX4 induces a transcriptome more characteristic of a less-differentiated cell state and inhibits myogenesis. J Cell Sci, 2016. 129(20): p. 3816–3831.

27. Bosnakovski, D., et al., DUX4c, an FSHD candidate gene, interferes with myogenic regulators and abolishes myoblast differentiation. Exp Neurol, 2008. 214(1): p. 87–96.

28. Bosnakovski, D., et al., An isogenetic myoblast expression screen identifies DUX4-mediated FSHD-associated molecular pathologies. EMBO J, 2008. 27(20): p. 2766–79.

29. Banerji, C.R.S., et al., PAX7 target genes are globally repressed in facioscapulohumeral muscular dystrophy skeletal muscle. Nat Commun, 2017. 8(1): p. 2152.

30. Banerji, C.R.S. and P.S. Zammit, PAX7 target gene repression is a superior FSHD biomarker than DUX4 target gene activation, associating with pathological severity and identifying FSHD at the single-cell level. Hum Mol Genet, 2019. 28(13): p. 2224–2236.

31. Rando, T.A., Role of nitric oxide in the pathogenesis of muscular dystrophies: a “two hit” hypothesis of the cause of muscle necrosis. Microsc Res Tech, 2001. 55(4): p. 223–35.

32. Tidball, J.G. and M. Wehling-Henricks, The role of free radicals in the pathophysiology of muscular dystrophy. J Appl Physiol (1985), 2007. 102(4): p. 1677–86.

33. Toscano, A., et al., Oxidative stress in myotonic dystrophy type 1. Free Radic Res, 2005. 39(7): p. 771–6.

34. Kramerova, I., et al., Mitochondrial abnormalities, energy deficit and oxidative stress are features of calpain 3 deficiency in skeletal muscle. Hum Mol Genet, 2009. 18(17): p. 3194–205.

35. Terrill, J.R., et al., Oxidative stress and pathology in muscular dystrophies: focus on protein thiol oxidation and dysferlinopathies. FEBS J, 2013. 280(17): p. 4149–64.

36. Winokur, S.T., et al., Facioscapulohumeral muscular dystrophy (FSHD) myoblasts demonstrate increased susceptibility to oxidative stress. Neuromuscul Disord, 2003. 13(4): p. 322–33.

37. Celegato, B., et al., Parallel protein and transcript profiles of FSHD patient muscles correlate to the D4Z4 arrangement and reveal a common impairment of slow to fast fibre differentiation and a general deregulation of MyoD-dependent genes. Proteomics, 2006. 6(19): p. 5303–21.

38. Cheli, S., et al., Expression profiling of FSHD-1 and FSHD-2 cells during myogenic differentiation evidences common and distinctive gene dysregulation patterns. PLoS One, 2011. 6(6): p. e20966.

39. Laoudj-Chenivesse, D., et al., Increased levels of adenine nucleotide translocator 1 protein and response to oxidative stress are early events in facioscapulohumeral muscular dystrophy muscle. J Mol Med (Berl), 2005. 83(3): p. 216–24.

40. Macaione, V., et al., RAGE-NF-kappaB pathway activation in response to oxidative stress in facioscapulohumeral muscular dystrophy. Acta Neurol Scand, 2007. 115(2): p. 115–21.

41. Sharma, V., et al., DUX4 differentially regulates transcriptomes of human rhabdomyosarcoma and mouse C2C12 cells. PLoS One, 2013. 8(5): p. e64691.

42. Turki, A., et al., Functional muscle impairment in facioscapulohumeral muscular dystrophy is correlated with oxidative stress and mitochondrial dysfunction. Free Radic Biol Med, 2012. 53(5): p. 1068–79.

43. Bou Saada, Y., et al., Facioscapulohumeral dystrophy myoblasts efficiently repair moderate levels of oxidative DNA damage. Histochem Cell Biol, 2016. 145(4): p. 475–83.

44. Dmitriev, P., et al., DUX4-induced constitutive DNA damage and oxidative stress contribute to aberrant differentiation of myoblasts from FSHD patients. Free Radic Biol Med, 2016. 99: p. 244–258.

45. Murphy, M.P., How mitochondria produce reactive oxygen species. Biochem J, 2009. 417(1): p. 1–13.

46. Thomas, L.W. and M. Ashcroft, Exploring the molecular interface between hypoxia-inducible factor signalling and mitochondria. Cell Mol Life Sci, 2019. 76(9): p. 1759–1777.

47. Le Moal, E., et al., Redox Control of Skeletal Muscle Regeneration. Antioxid Redox Signal, 2017. 27(5): p. 276–310.

48. Banerji, C.R.S., et al., Dynamic transcriptomic analysis reveals suppression of PGC1alpha/ERRalpha drives perturbed myogenesis in facioscapulohumeral muscular dystrophy. Hum Mol Genet, 2019. 28(8): p. 1244–1259.

49. Barro, M., et al., Myoblasts from affected and non-affected FSHD muscles exhibit morphological differentiation defects. J Cell Mol Med, 2010. 14(1-2): p. 275–89.

50. Banerji, C.R., et al., beta-Catenin is central to DUX4-driven network rewiring in facioscapulohumeral muscular dystrophy. J R Soc Interface, 2015. 12(102): p. 20140797.

51. Lek, A., et al., Applying genome-wide CRISPR-Cas9 screens for therapeutic discovery in facioscapulohumeral muscular dystrophy. Sci Transl Med, 2020. 12(536).

52. Denny, A.P. and A.K. Heather, Are Antioxidants a Potential Therapy for FSHD? A Review of the Literature. Oxid Med Cell Longev, 2017. 2017: p. 7020295.

53. Bosnakovski, D., et al., High-throughput screening identifies inhibitors of DUX4-induced myoblast toxicity. Skelet Muscle, 2014. 4(1): p. 4.

54. Sasaki-Honda, M., et al., A patient-derived iPSC model revealed oxidative stress increases facioscapulohumeral muscular dystrophy-causative DUX4. Hum Mol Genet, 2018. 27(23): p. 4024–4035.

55. Emilie, P., et al., Oxidative stress and dystrophy Facioscapulohumeral: Effects of vitamin C, vitamin E, zinc gluconate and selenomethionine supplementation. Free Radic Biol Med, 2014. 75 Suppl 1: p. S14.

56. Passerieux, E., et al., Effects of vitamin C, vitamin E, zinc gluconate, and selenomethionine supplementation on muscle function and oxidative stress biomarkers in patients with facioscapulohumeral dystrophy: a double-blind randomized controlled clinical trial. Free Radic Biol Med, 2015. 81: p. 158–69.

57. Karpukhina, A., et al., Analysis of genes regulated by DUX4 via oxidative stress reveals potential therapeutic targets for treatment of facioscapulohumeral dystrophy. Redox Biology, 2021. 43: p. 102008.

58. Wang, L.H., et al., MRI-informed muscle biopsies correlate MRI with pathology and DUX4 target gene expression in FSHD. Hum Mol Genet, 2019. 28(3): p. 476–486.

59. Zorova, L.D., et al., Mitochondrial membrane potential. Anal Biochem, 2018. 552: p. 50–59.

60. Korshunov, S.S., V.P. Skulachev, and A.A. Starkov, High protonic potential actuates a mechanism of production of reactive oxygen species in mitochondria. FEBS Lett, 1997. 416(1): p. 15–8.

61. Jagannathan, S., et al., Model systems of DUX4 expression recapitulate the transcriptional profile of FSHD cells. Hum Mol Genet, 2016. 25(20): p. 4419–4431.

62. Bhattacharya, D. and A. Scime, Mitochondrial Function in Muscle Stem Cell Fates. Front Cell Dev Biol, 2020. 8: p. 480.

63. Haramizu, S., et al., Dietary resveratrol confers apoptotic resistance to oxidative stress in myoblasts. J Nutr Biochem, 2017. 50: p. 103–115.

64. Olah, G., et al., Differentiation-Associated Downregulation of Poly(ADP-Ribose) Polymerase-1 Expression in Myoblasts Serves to Increase Their Resistance to Oxidative Stress. PLoS One, 2015. 10(7): p. e0134227.

65. Hagen, T., et al., Redistribution of intracellular oxygen in hypoxia by nitric oxide: effect on HIF1alpha. Science, 2003. 302(5652): p. 1975–8.

66. Hamanaka, R.B. and N.S. Chandel, Mitochondrial reactive oxygen species regulate hypoxic signaling. Curr Opin Cell Biol, 2009. 21(6): p. 894–9.

67. Solaini, G., et al., Hypoxia and mitochondrial oxidative metabolism. Biochim Biophys Acta, 2010. 1797(6-7): p. 1171–7.

68. Pala, F., et al., Distinct metabolic states govern skeletal muscle stem cell fates during prenatal and postnatal myogenesis. J Cell Sci, 2018. 131(14).

69. Sin, J., et al., Mitophagy is required for mitochondrial biogenesis and myogenic differentiation of C2C12 myoblasts. Autophagy, 2016. 12(2): p. 369–80.

70. Magalhaes, J., et al., Acute and severe hypobaric hypoxia increases oxidative stress and impairs mitochondrial function in mouse skeletal muscle. J Appl Physiol (1985), 2005. 99(4): p. 1247–53.

71. Bittel, A.J., et al., Membrane Repair Deficit in Facioscapulohumeral Muscular Dystrophy. Int J Mol Sci, 2020. 21(15).

72. Tassin, A., et al., DUX4 expression in FSHD muscle cells: how could such a rare protein cause a myopathy? J Cell Mol Med, 2013. 17(1): p. 76–89.

73. Suski, J.M., et al., Relation between mitochondrial membrane potential and ROS formation. Methods Mol Biol, 2012. 810: p. 183–205.

74. Radi, R., Oxygen radicals, nitric oxide, and peroxynitrite: Redox pathways in molecular medicine. Proc Natl Acad Sci U S A, 2018. 115(23): p. 5839–5848.

75. Bohnert, K.R., J.D. McMillan, and A. Kumar, Emerging roles of ER stress and unfolded protein response pathways in skeletal muscle health and disease. J Cell Physiol, 2018. 233(1): p. 67–78.

76. Homma, S., et al., Expression of FSHD-related DUX4-FL alters proteostasis and induces TDP-43 aggregation. Ann Clin Transl Neurol, 2015. 2(2): p. 151–66.

77. Pirie, E., et al., S-nitrosylated TDP-43 triggers aggregation, cell-to-cell spread, and neurotoxicity in hiPSCs and in vivo models of ALS/FTD. Proc Natl Acad Sci U S A, 2021. 118(11).

78. Jagannathan, S., et al., Quantitative proteomics reveals key roles for post-transcriptional gene regulation in the molecular pathology of facioscapulohumeral muscular dystrophy. Elife, 2019. 8.

79. Machado, J.D., et al., Nitric oxide modulates a late step of exocytosis. J Biol Chem, 2000. 275(27): p. 20274–9.

80. Matsushita, K., et al., Nitric oxide regulates exocytosis by S-nitrosylation of N-ethylmaleimide-sensitive factor. Cell, 2003. 115(2): p. 139–50.

81. Brown, G.C., Nitric oxide and mitochondrial respiration. Biochim Biophys Acta, 1999. 1411(2-3): p. 351–69.

82. Brown, G.C. and V. Borutaite, Inhibition of mitochondrial respiratory complex I by nitric oxide, peroxynitrite and S-nitrosothiols. Biochim Biophys Acta, 2004. 1658(1-2): p. 44–9.

83. Erusalimsky, J.D. and S. Moncada, Nitric oxide and mitochondrial signaling: from physiology to pathophysiology. Arterioscler Thromb Vasc Biol, 2007. 27(12): p. 2524–31.

84. Galkin, A., A. Higgs, and S. Moncada, Nitric oxide and hypoxia. Essays Biochem, 2007. 43: p. 29–42.

85. Taylor, C.T. and S. Moncada, Nitric oxide, cytochrome C oxidase, and the cellular response to hypoxia. Arterioscler Thromb Vasc Biol, 2010. 30(4): p. 643–7.

86. Brune, B., Nitric oxide: NO apoptosis or turning it ON? Cell Death Differ, 2003. 10(8): p. 864–9.

87. Kim, P.K., et al., The regulatory role of nitric oxide in apoptosis. Int Immunopharmacol, 2001. 1(8): p. 1421–41.

88. Kussmaul, L. and J. Hirst, The mechanism of superoxide production by NADH:ubiquinone oxidoreductase (complex I) from bovine heart mitochondria. Proc Natl Acad Sci U S A, 2006. 103(20): p. 7607–12.

89. Brand, M.D., The sites and topology of mitochondrial superoxide production. Exp Gerontol, 2010. 45(7-8): p. 466–72.

90. Forkink, M., et al., Mitochondrial hyperpolarization during chronic complex I inhibition is sustained by low activity of complex II, III, IV and V. Biochim Biophys Acta, 2014. 1837(8): p. 1247–56.

91. Giovannini, C., et al., Mitochondria hyperpolarization is an early event in oxidized low-density lipoprotein-induced apoptosis in Caco-2 intestinal cells. FEBS Lett, 2002. 523(1-3): p. 200–6.

92. Piacentini, M., et al., Transglutaminase overexpression sensitizes neuronal cell lines to apoptosis by increasing mitochondrial membrane potential and cellular oxidative stress. J Neurochem, 2002. 81(5): p. 1061–72.

93. Vyssokikh, M.Y., et al., Mild depolarization of the inner mitochondrial membrane is a crucial component of an anti-aging program. Proc Natl Acad Sci U S A, 2020. 117(12): p. 6491–6501.

94. Lambert, A.J. and M.D. Brand, Superoxide production by NADH:ubiquinone oxidoreductase (complex I) depends on the pH gradient across the mitochondrial inner membrane. Biochem J, 2004. 382(Pt 2): p. 511–7.

95. Chinopoulos, C. and V. Adam-Vizi, Mitochondria as ATP consumers in cellular pathology. Biochim Biophys Acta, 2010. 1802(1): p. 221–7.

96. Xu, W., I.G. Charles, and S. Moncada, Nitric oxide: orchestrating hypoxia regulation through mitochondrial respiration and the endoplasmic reticulum stress response. Cell Res, 2005. 15(1): p. 63–5.

97. Wagatsuma, A. and K. Sakuma, Mitochondria as a potential regulator of myogenesis. ScientificWorldJournal, 2013. 2013: p. 593267.

98. Clanton, T.L., Hypoxia-induced reactive oxygen species formation in skeletal muscle. J Appl Physiol (1985), 2007. 102(6): p. 2379–88.

99. Waypa, G.B., et al., Increases in mitochondrial reactive oxygen species trigger hypoxia-induced calcium responses in pulmonary artery smooth muscle cells. Circ Res, 2006. 99(9): p. 970–8.

100. Chandel, N.S., et al., Mitochondrial reactive oxygen species trigger hypoxia-induced transcription. Proc Natl Acad Sci U S A, 1998. 95(20): p. 11715–20.

101. Chandel, N.S., et al., Reactive oxygen species generated at mitochondrial complex III stabilize hypoxia-inducible factor-1alpha during hypoxia: a mechanism of O2 sensing. J Biol Chem, 2000. 275(33): p. 25130–8.

102. van der Kooi, E.L., et al., No effect of folic acid and methionine supplementation on D4Z4 methylation in patients with facioscapulohumeral muscular dystrophy. Neuromuscul Disord, 2006. 16(11): p. 766–9.

103. Murphy, M.P., Antioxidants as therapies: can we improve on nature? Free Radic Biol Med, 2014. 66: p. 20–3.

104. Snow, B.J., et al., A double-blind, placebo-controlled study to assess the mitochondria-targeted antioxidant MitoQ as a disease-modifying therapy in Parkinson’s disease. Mov Disord, 2010. 25(11): p. 1670–4.

105. Gane, E.J., et al., The mitochondria-targeted anti-oxidant mitoquinone decreases liver damage in a phase II study of hepatitis C patients. Liver Int, 2010. 30(7): p. 1019–26.

106. Petrov, A., et al., SkQ1 Ophthalmic Solution for Dry Eye Treatment: Results of a Phase 2 Safety and Efficacy Clinical Study in the Environment and During Challenge in the Controlled Adverse Environment Model. Adv Ther, 2016. 33(1): p. 96–115.

107. E, B.D. and G. Marfany, The Relevance of Oxidative Stress in the Pathogenesis and Therapy of Retinal Dystrophies. Antioxidants (Basel), 2020. 9(4).

108. Forli, F., et al., Mitochondrial syndromic sensorineural hearing loss. Biosci Rep, 2007. 27(1-3): p. 113–23.

109. Kokotas, H., M.B. Petersen, and P.J. Willems, Mitochondrial deafness. Clin Genet, 2007. 71(5): p. 379–91.

110. Tavanai, E. and G. Mohammadkhani, Role of antioxidants in prevention of age-related hearing loss: a review of literature. European Archives of Oto-Rhino-Laryngology, 2017. 274(4): p. 1821–1834.

111. Fujimoto, C. and T. Yamasoba, Mitochondria-Targeted Antioxidants for Treatment of Hearing Loss: A Systematic Review. Antioxidants (Basel), 2019. 8(4).

112. Jiang, Q., et al., Mitochondria-Targeted Antioxidants: A Step towards Disease Treatment. Oxid Med Cell Longev, 2020. 2020: p. 8837893.

113. Oyewole, A.O. and M.A. Birch-Machin, Mitochondria-targeted antioxidants. FASEB J, 2015. 29(12): p. 4766–71.

114. Badodi, S., et al., Inositol treatment inhibits medulloblastoma through suppression of epigenetic-driven metabolic adaptation. Nat Commun, 2021. 12(1): p. 2148.

115. Tarazona, S., et al., Data quality aware analysis of differential expression in RNA-seq with NOISeq R/Bioc package. Nucleic Acids Res, 2015. 43(21): p. e140.

116. Durinck, S., et al., Mapping identifiers for the integration of genomic datasets with the R/Bioconductor package biomaRt. Nat Protoc, 2009. 4(8): p. 1184–91.

117. Robinson, M.D., D.J. McCarthy, and G.K. Smyth, edgeR: a Bioconductor package for differential expression analysis of digital gene expression data. Bioinformatics, 2010. 26(1): p. 139–40.

118. Shannon, P., et al., Cytoscape: a software environment for integrated models of biomolecular interaction networks. Genome Res, 2003. 13(11): p. 2498–504.

119. Krom, Y.D., et al., Generation of isogenic D4Z4 contracted and noncontracted immortal muscle cell clones from a mosaic patient: a cellular model for FSHD. Am J Pathol, 2012. 181(4): p. 1387–401.

120. Homma, S., et al., A unique library of myogenic cells from facioscapulohumeral muscular dystrophy subjects and unaffected relatives: family, disease and cell function. Eur J Hum Genet, 2012. 20(4): p. 404–10.

121. Choi, S.H., et al., DUX4 recruits p300/CBP through its C-terminus and induces global H3K27 acetylation changes. Nucleic Acids Res, 2016. 44(11): p. 5161–73.

122. Rosca, M.G., et al., Cardiac mitochondria in heart failure: decrease in respirasomes and oxidative phosphorylation. Cardiovasc Res, 2008. 80(1): p. 30–9.

