## Supplemental Figures for "Interplay between mitochondria and reactive oxygen and nitrogen species in metabolic adaptation to hypoxia in facioscapulohumeral muscular dystrophy: potential therapeutic targets"

### SUPPLEMENTARY MATERIAL:

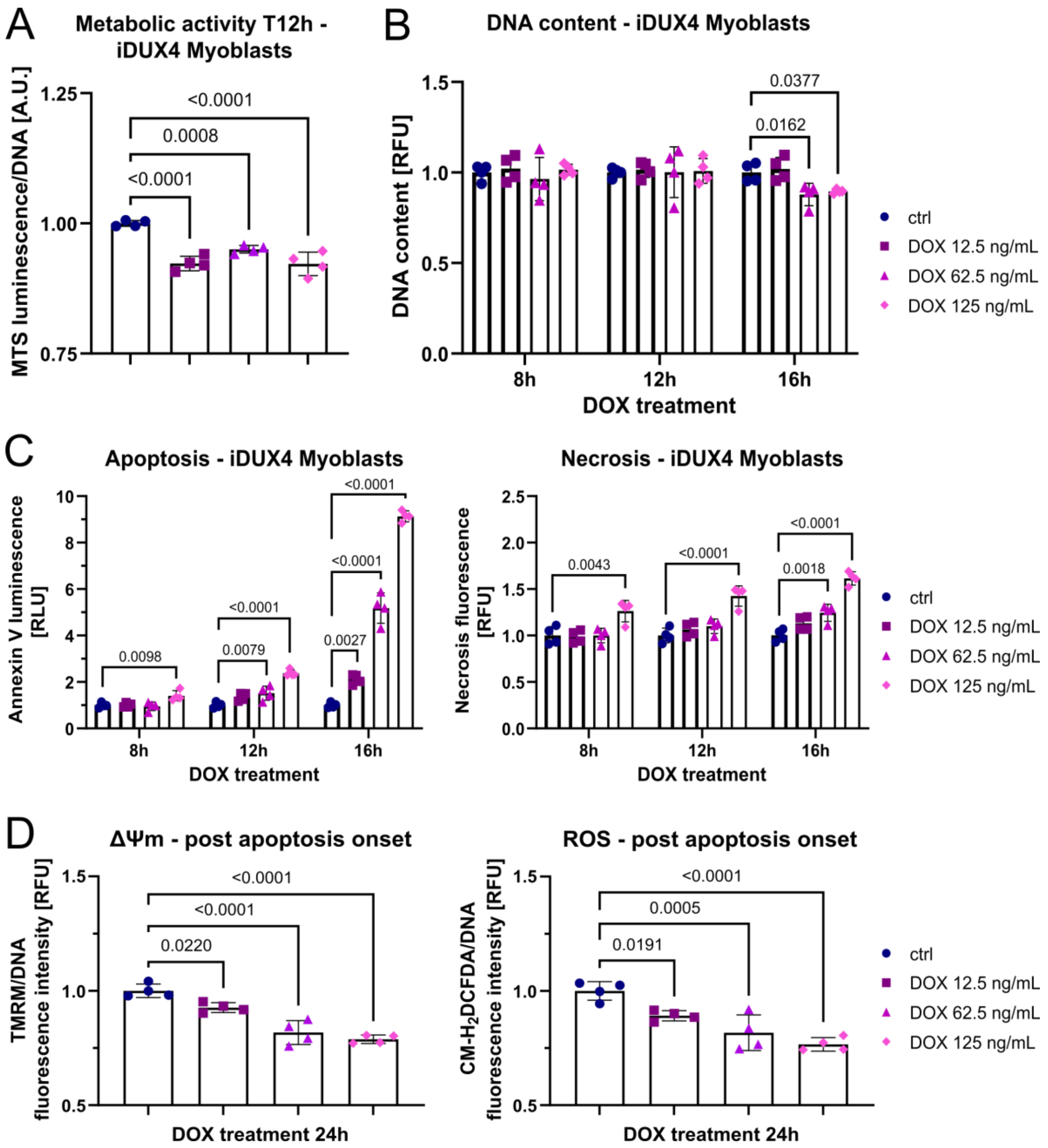


**Fig. S1:** *Changes in metabolic activity precede apoptosis.*

**(A)** Metabolic activity is significantly reduced after 12h of variable DUX4 expression in iDUX4 myoblasts, before elevated ROS levels can be observed. **(B)** DNA content in iDUX4 myoblasts subjected to variable DUX4 expression for up to 16h shows decrease in cell viability at the 16h timepoint. **(C)** Dose-dependent effect of DUX4 on apoptosis and necrosis in iDUX4 myoblasts subjected to variable DUX4 expression over time revealing apoptosis as major driver of DUX4-induced cell death: onset of apoptosis at around 12h, with significant increase of apoptosis and necrosis in all DOX treatment groups after 16h. **(D)** ΔΨm and ROS levels 24h after variable DUX4 expression in myoblasts ("post-apoptosis onset") demonstrating collapse of ΔΨm with concomitant decrease of ROS levels due to mitochondria-mediated death. Data is mean ± s.d. and normalised to untreated controls for each timepoint, from 3-4 wells each from a representative experiment with *p* values as indicated.

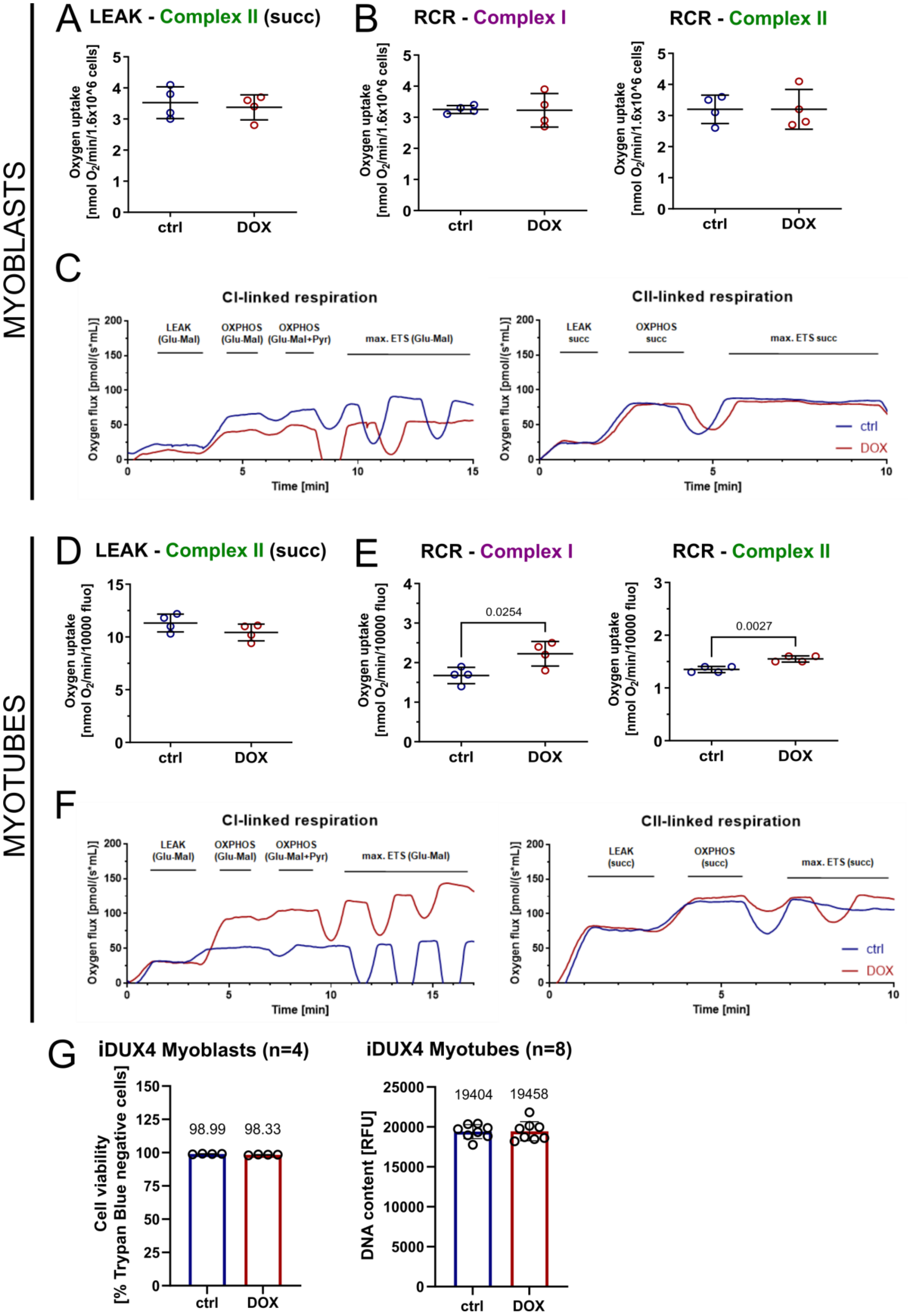


**Fig. S2:** *Effects of DUX4 expression on mitochondrial respiration.*

**(A, B)** LEAK (uncoupled) respiration through complex II is not affected by DUX4 expression (DOX 62.5 ng/mL for 16h) in iDUX4 myoblasts, neither are the respiratory control ratios (RCR) through complex I and II. **(C)** Representative oxygraphs for high-resolution respirometry experiments in iDUX4 myoblasts (controls versus DUX4 expression): complex I- and complex II-linked respiration. **(D, E)** LEAK (uncoupled) respiration through complex II is not affected by DUX4 expression (DOX 62.5 ng/mL for 24h) in iDUX4 myotubes, but respiratory control ratios RCR through complex I and II are slightly elevated. **(F)** Representative oxygraphs for high-resolution respirometry experiments in myotubes (controls versus DUX4 expression): complex I- and complex II-linked respiration. **(G)** Results of cell viability (for iDUX4 myoblasts; left graph) and DNA quantitation (for iDUX4 myotubes; right graph) measurements of cells prior to respirometry for normalisation of $O_2$ uptake. Data is mean ± s.d. from 4-8 wells each from a representative experiment with *p* values as indicated.

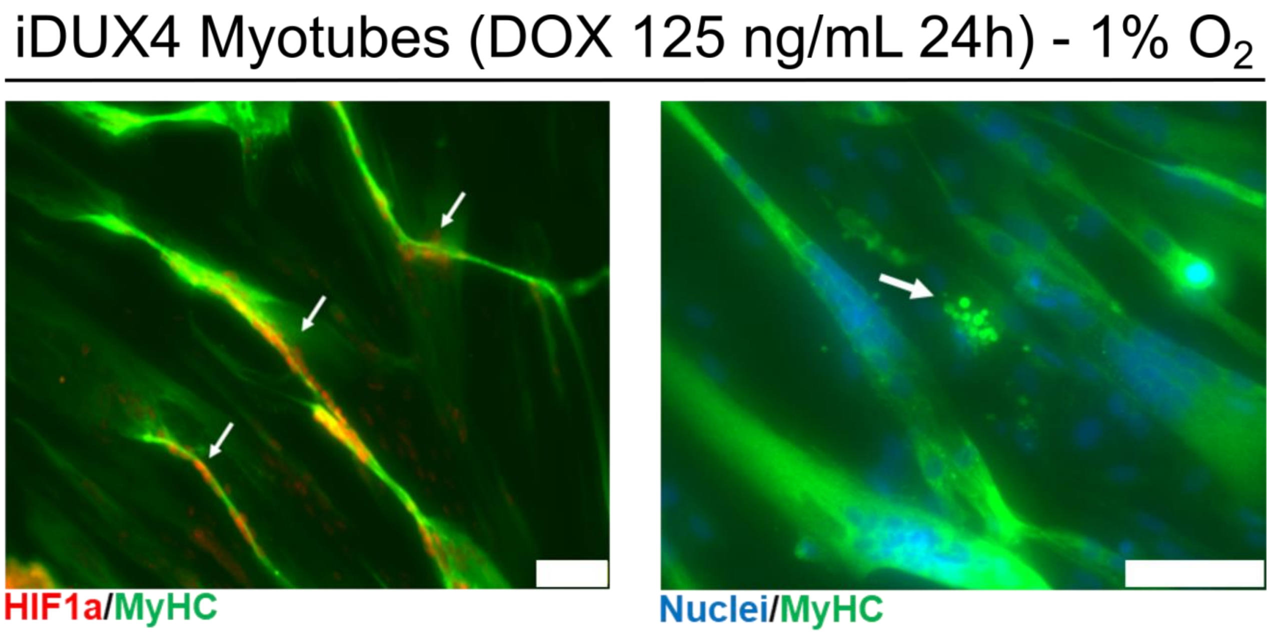


**Fig. S3:** *Morphological changes in response to DUX4 expression in hypoxic myotubes.* Immunofluorescence micrographs from iDUX4 myotubes expressing DUX4 (DOX 125 ng/mL for 24) in hypoxia. Left: strong enrichment of HIF1α-postive myonuclei (marked by arrows) in hypotrophic/atrophic myotubes (scale bar represents 75 μm). Right: myotube fragmentation (marked by arrow) indicative of myotube death (scale bar represents 50 μm).
